# Structural basis of calcium-mediated conformational dynamics and cargo binding in Vacuolar Sorting Receptor 1 (VSR1)

**DOI:** 10.1101/2024.12.07.627339

**Authors:** Thomas H. Wilson, Rebecca J. Parr, Rafael J. Borges, Mahmoud K. Eldahshoury, Nita Shah, Joanne E. Nettleship, Marilyn Paul, Paul Fyfe, Jing An, Line G Kristensen, Sayan Gupta, Kamel El Omari, James Birch, Ana J. Loureiro Levada, Marcos R. M. Fontes, Jurgen Denecke, Raymond J. Owens, Corie Y Ralston, Adrian Goldman, Vincent Postis, Isabel Usón, Andrew Quigley, Carine de Marcos Lousa

**Author notes:** These authors contributed equally to this work (Thomas H. Wilson, Rebecca J. Parr, Rafael J. Borges). NS: Evotec (United Kingdom), Innovation Drive, Milton Park, Abingdon OX14 4RZ, United Kingdom. MP: Biophysics Facility, Department of Biochemistry, University of Cambridge.

## Abstract

Vacuolar sorting receptors (VSRs) are type I membrane proteins crucial for seed germination and plant development. Although VSR trafficking has been extensively studied, the mechanism of cargo recognition and release by the N-terminal lumenal region remains poorly understood. Here, we solved the crystal structures of both unloaded and cargo-loaded forms of the VSR1 lumenal region comprising the protease-associated (PA) domain, the Central domain and epidermal growth factor-like repeats. Two cargo binding sites were identified: one previously reported site found in the PA and Central domains and one uncharacterised site located within the Central domain. The large conformational differences observed between the two crystal structures are confirmed in solution by Small-angle X-ray scattering (SAXS) experiments. Their combination revealed that calcium coordination remodels inter-domain linkers, triggering large conformational changes that expose and remodel the two cargo binding sites. Complementary X-ray footprinting mass spectrometry (XFMS) mapped the residues undergoing solvent accessibility changes induced by cargo binding and confirmed residues affected in both cargo binding sites and across all three domains in solution. Together, these findings reveal a dynamic calcium-dependent mechanism in which high calcium concentrations promote cargo binding, while cargo release is favoured by lower calcium concentrations and pH. This work provides a structural framework for understanding VSR-mediated vacuolar trafficking and cargo sorting in plants.

## Introduction

Vacuolar Sorting Receptors (VSRs) are type I transmembrane proteins that mediate the delivery of soluble cargo proteins to the plant vacuole, a process essential for seed germination, protein storage, cellular homeostasis and plant development. Following cargo capture in early secretory compartments, VSRs traffic through the endomembrane system and, after cargo release, subsequently recycle to early compartments. *Arabidopsis thaliana* contains seven VSR isoforms, all of which are composed of three regions: the N-terminal lumenal region, a transmembrane region and C-terminal region which contains several sorting signals involved in trafficking^1–7^. Cargo recognition is mediated by the lumenal region which consists of a protease-associated (PA) domain, a Central domain, and three epidermal growth factor-like (EGF) repeats^8–11^. VSRs recognise a variety of cargo proteins carrying either a sequence-specific determinant (ssVSD) or a C-terminal determinant (ctVSD). While ssVSD are typically located at the N-terminus and contain a signal motif which sequence can vary around the well characterised NPIR peptide motif ^8–16^, ctVSDs lack a defined consensus sequence but are characterised by hydrophobic residues at the C-terminus^17–19^. Despite extensive characterisation of cargo motifs and receptor trafficking pathways, the molecular basis of cargo recognition and release by VSRs remains poorly understood.

Several factors have been implicated in modulating VSR function. Calcium binding by the EGF repeats appear to stabilise receptor-cargo interactions^15^, while N-linked glycosylation promotes cargo binding, without affecting trafficking^20^. In addition, VSR characteristically contains a high proportion of cysteine residues (i.e. VSR1 has 35 cysteines in its lumenal domain), which were recently postulated to play a role in stabilising vacuolar cargo through oxidative reactions^21^. Collectively, these observations indicate that cargo recognition involves coordinated structural regulation, yet the underlying mechanism linking these factors to cargo loading and release remains unclear.

Previously, the bacterially-expressed PA domain of *A. thaliana* VSR1 was crystallized in both unbound and complexed form, with either ssVSD or ctVSD small peptide. The structures revealed the presence of a putative cargo-binding site centred around Arg95 interactions, as well as local structural changes of switch regions upon cargo binding, but lacks the context of the remaining two domains^10,11,22^. In addition, the signal motif of the ssVSD peptide was not well resolved. Recently, based on the analysis of original diffraction data collected in 2004, the unloaded form of the VSR1 lumenal region containing the three sub-domains expressed in insect cells was reported to crystalise as a trimer (PDB id: 9B1R) ^21, 23^, but the cargo loaded form was not reported.

Clearly, further structural information on the full N-terminal region of VSR bound to vacuolar cargo is needed to enlighten the mechanism of receptor-cargo recognition and release.

Here, we combined X-ray crystallography, small-angle X-ray scattering (SAXS) and X-ray footprinting mass spectrometry (XFMS) to investigate the structural basis of cargo recognition by VSR1. We determined two structures of the VSR1 lumenal region: one unloaded and one cargo-loaded form with a classical ssVSD peptide. SAXS demonstrated the role of calcium and pH in modulating the conformation of the overall VSR1 lumenal region and its cargo binding capabilities. In parallel, XFMS mapped changes in solvent accessibility of residues located throughout the receptor and induced by cargo recognition. Together, these findings support a dynamic model in which calcium-dependent structural remodelling regulates cargo loading and release, providing a molecular framework for the understanding of VSR-mediated vacuolar trafficking.

## Results

### Unloaded VSR1-N_t_ structure contains predicted glycosylations and adopts a trimeric assembly

The N-terminal domain of VSR1 (VSR1-Nt) containing the PA domain, the Central domain and the three EGF repeats, was expressed and purified from Expi293 cells (**SI Fig. 1**). Crystals belonged to the P2_1_3 space group and diffracted to a resolution limit of 3.5 Å (**SI Table 1**). Structure determination required the development of an ARCIMBOLDO-based phasing strategy, as only the isolated PA domain structure was available for molecular replacement at the time of analysis^24–29^.

**Figure 1.**
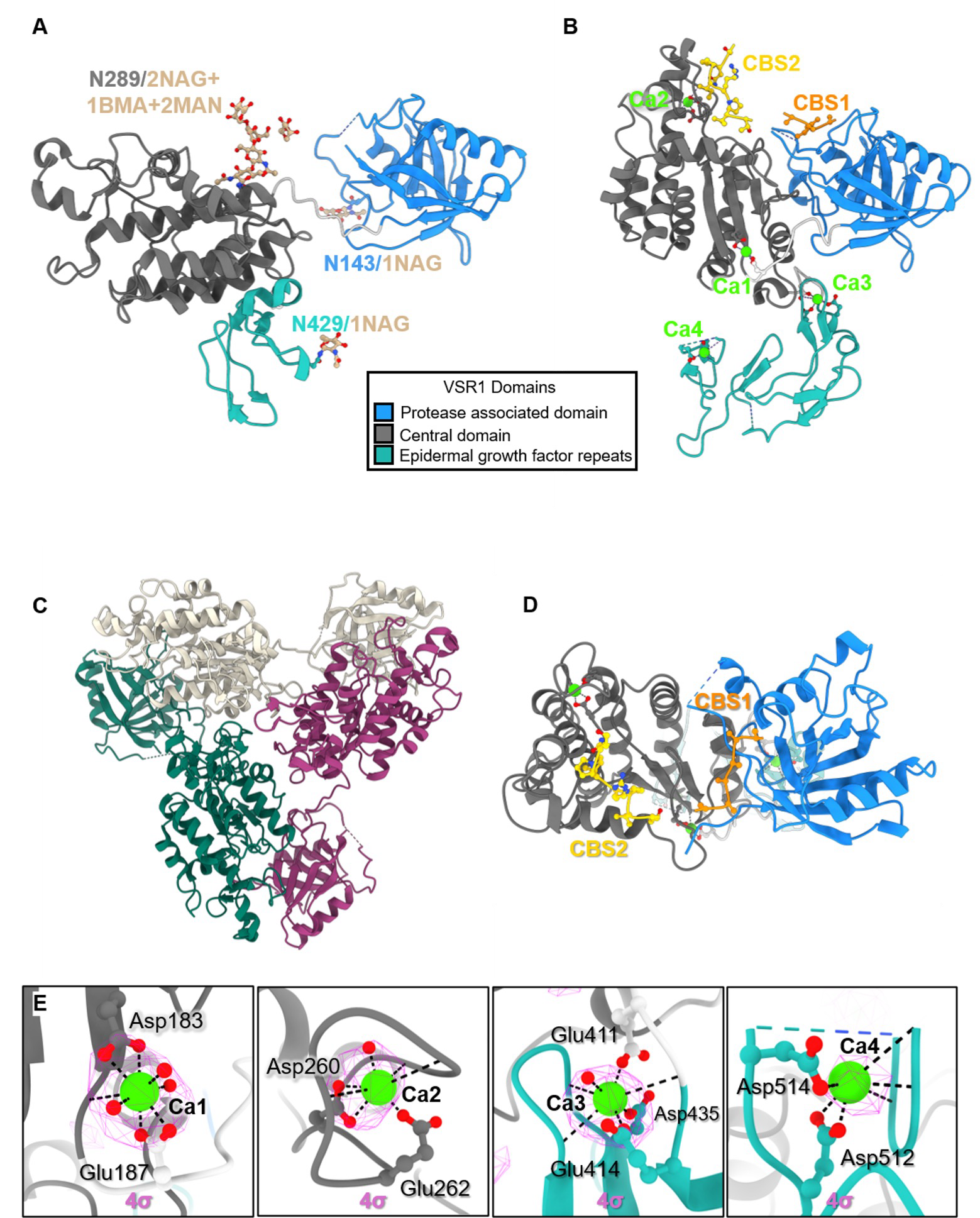
The crystal structure of unloaded and cargo loaded VSR1-N_t_ lumenal domain. The structure of VSR containing the three domains: Protease associated domain (light blue), Central domain (black) and EGF repeats (green). Absent residues in the PA main chain are shown in dashed lines. **A. Monomeric unloaded structure.** Glycosylations are shown in yellow at the three sites (N143 in the PA domain, N289 in the Central domain and N249 in the first epidermal growth factor-like (EGF) repeat). **B. Top view of the trimer of the unloaded structure**: crystal structure showing the arrangement of the three monomers A, B and C. B and C are symmetry equivalents of Chain A. **C. Monomeric cargo-loaded structure.** The peptides are represented in orange for cargo binding site 1 (CBS1) and yellow for Cargo binding site 2 (CBS2). The four Calcium ions are shown in bright green and names Ca1 to Ca4. **D. Top view of the monomeric cargo-loaded structure**. The peptides are represented in orange for cargo binding site 1 (CBS1) and yellow for Cargo binding site 2 (CBS2). **E.** Detail of the four calcium coordination sites with anomalous maps at Calcium edge in pink and residues and water in sticks.

The resulting model comprised residues 20-462 and contains the PA domain, Central domain and one EGF domain (**Fig. 1A**). The overall structure adopts a trimeric assembly with C3 symmetry (**Fig. 1C**), consistent with that reported by Park *et al*.,^21^. In our structure, N-linked glycosylation was observed at three sites, Asn143, Asn289 and Asn429 (**Fig. 1A and SI Fig.2**). Electron density at Asn289 allowed modelling of a 2NAG–1BMA–2MAN glycan, whereas only the first NAG could be modelled at Asn143 and Asn429 ^26^. The structure contains 10 disulfide bridges: one in the PA domain, six in the Central domain and three in the first EGF repeat, stabilizing the overall structure of VSR1-N_t_ **(SI Table 2**). A single calcium ion was identified by anomalous diffraction; however, it is located on a crystallographic threefold axis near Asp344 and is unlikely to represent a biologically relevant calcium-binding site.

**Figure 2.**
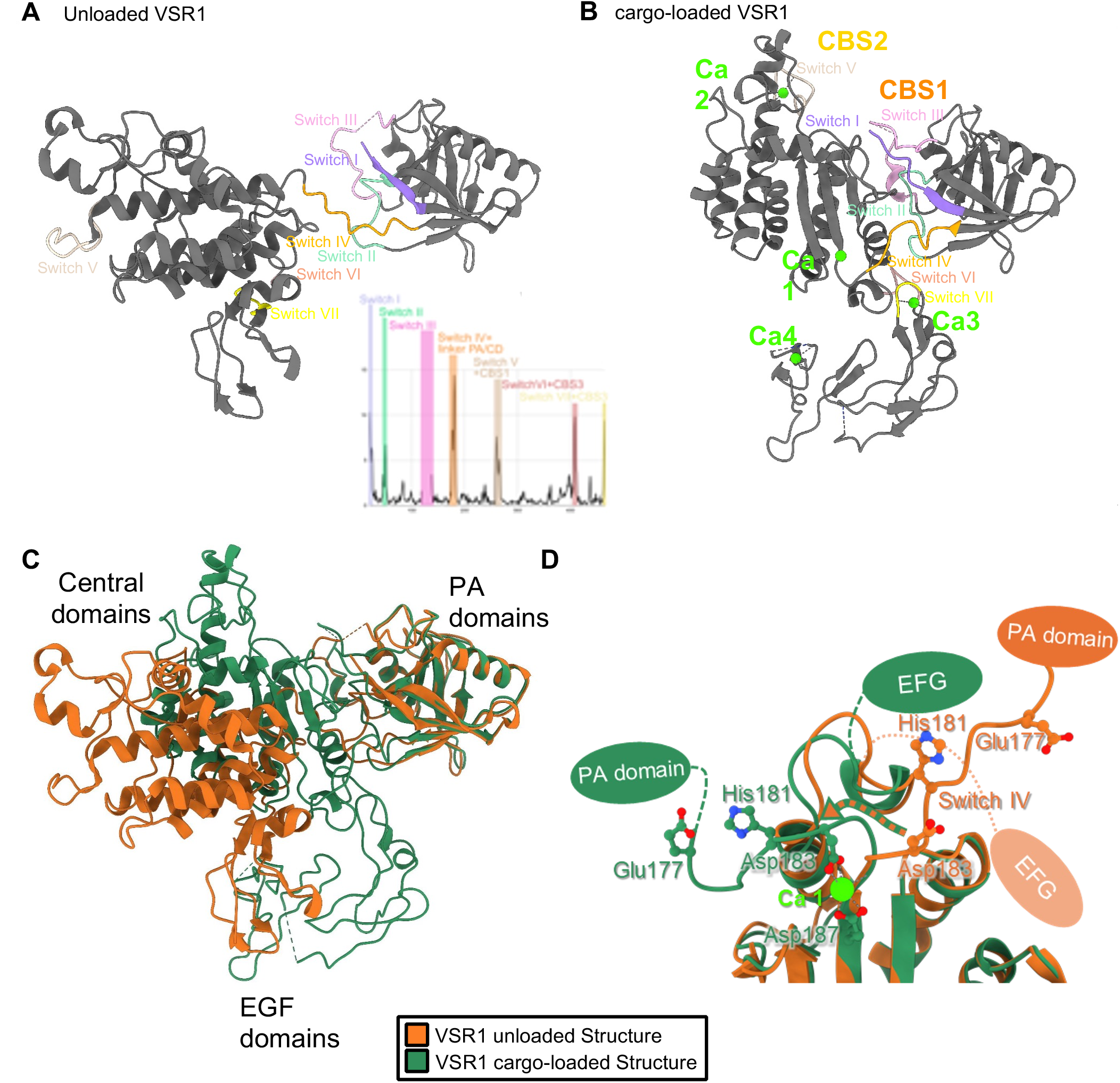
The rearrangements between the unloaded and peptide cargo-loaded form of VSR1-N_t_. **A.** The structure of the unloaded VSR1-N_t_ monomer showing the position of switches I-IV (Luo *et al.,* 2014) and new switches V to VII. The insert plots values of Cα displacement comparing the unloaded and cargo-loaded structures, large values showing the seven switches are highlighted in different colours **B.** The structure of the cargo-loaded VSR1-N_t_ showing the position of switches I-IV (Luo *et al*., 2014) and new switches V to VII. Peptides in cargo binding site 1 and 2 are represented in yellow and orange lines and the four coordinated calcium ions are represented in bright green spheres. **C.** Zoom onto switch IV differences between unloaded (orange) and cargo-loaded (dark green) structures. **D.** Zoom in the switch IV. Calcium 1 ( Ca1 light green sphere) coordination by side chains of D183 and E187 induces large domain movements (from orange to green) concomitant with a profound remodelling of switch IV stabilised by a new salt bridge formed between H181 with E177. Wilson et al., 2026

### Cargo-loaded VSR1-N_t_ crystal structure reveals two cargo binding sites (CBS) and 4 calcium ions bound

Co-crystallisation with an Aleurain-GFP recombinant protein was unsuccessful. However, co-crystallisation with the barley (Hordeum Vulgare) ssVSD peptide (1ADSNPIRPVT10), previously used by Lu *et al.* ^10^, yielded crystals. These belonged to space group I222, distinct from the unloaded structure, that diffracted to a resolution limit of 1.87 Å (**SI. Table 1**). The cargo-bound VSR1-Nt structure crystallised as a monomer and comprise the PA domain, central domain and two EGF domains, with the third EGF domain unresolved (**Fig. 1B**).

Electron density corresponding to bound peptides was observed at two distinct sites, named cargo binding sites CBS1 and CBS2 (**Fig. 1D**). CBS1 corresponds to a previously reported site involving Arg95^10^, whereas CBS2 represents a novel uncharacterised cargo-binding site. A partially resolved N-linked glycan could be modelled at Asn289 allowing modelling of 2NAG-1BMA-1MAN glycan. However, no density corresponding to the glycosylation was observed at Asn143 or Asn429 in the cargo-loaded structure (**SI Fig.2**).

Four calcium-binding sites were identified within the cargo-bound structure (**Fig. 1E**). Their assignment was independently verified by calcium anomalous diffraction, with four anomalous peaks observed at the calcium absorption edge (λ = 3.07 Å) that disappeared below the edge (λ = 3.09 Å) (**SI Table 4**). One calcium ion is located at the PA–central domain interface (Ca1), a second within the central domain (Ca2), and the remaining two within the first and second EGF domains (Ca3, Ca4), respectively. These observations demonstrate extensive calcium coordination throughout the cargo-bound VSR1-Nt structure.

### Three additional switches identified undergo wide conformation changes which contribute to cargo binding and calcium binding sites formation

The unloaded and cargo loaded structures were compared to identify main rearrangements occurring upon calcium coordination and cargo binding (**Fig. 2**). Four conformational switches (I to IV) were previously described in the PA domain^10^ and confirmed here to undergo large rearrangements.

The new cargo-loaded form presented here reveals three additional switches undergoing large conformational changes: switch V (259-268) in the Central domain, switch VI (407-412) and switch VII (436-438) in EGF domains (**Fig. 2A, 2B, and SI Table 2**). In particular, switch V and VII contain residues involved in calcium coordination at Ca2 and Ca3 sites respectively.

Upon calcium binding, switch I and switch III, which were partially disordered in the unloaded form, undergo a conformational change to form part of CBS1 (see below). Switch II (especially _51_QY_52_), which was engaged in an interaction with the CD of the opposite monomer in the trimer of the unloaded form, now stabilizes the helix (385-395) between the CD and the first EGF repeat in the cargo loaded form. Switch IV undergoes a large remodelling, facilitated by the Ca1 coordination described above and stabilised by the newly formed bridge between Glu177-His181, moving the CD and the PA domain closer (**Fig. 2C and 2D**). Switch V (Central domain), which was interacting with switch I (PA domain) of the opposite monomer in the trimer, is now positioned close to cargo binding site 2 (CBS2) and Ca2 in the loaded form (**Fig. 2B**). Finally, switch VI and VII undergo a rearrangement, bringing the EGF domain closer to the PA domain.

Altogether, the large conformational changes observed upon calcium coordination result in the formation and exposure of the two cargo binding sites, which were hidden in the unloaded trimeric structure (**SI Fig. 3**).

**Figure 3.**
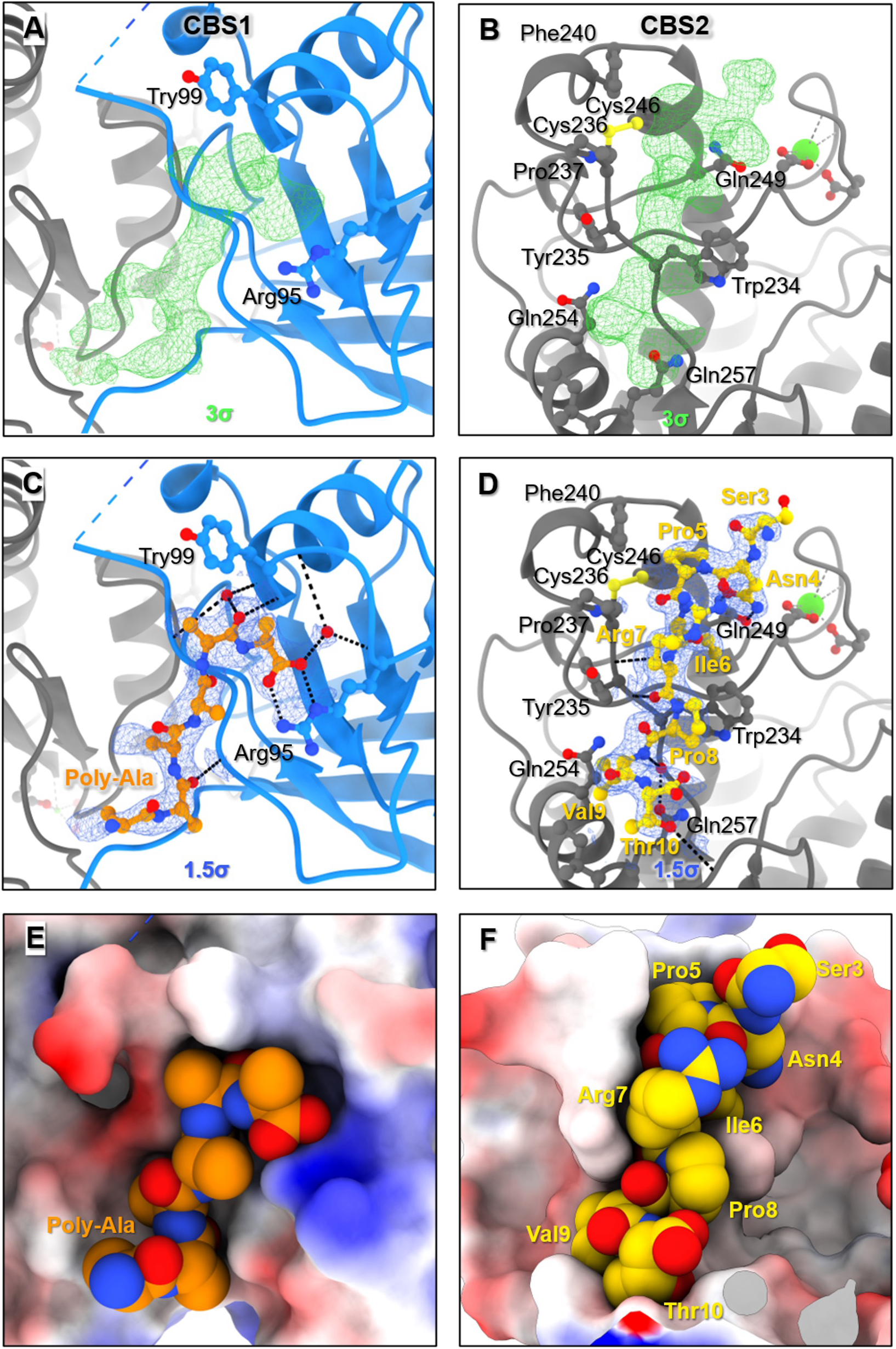
The two peptide cargo-binding sites (CBS) in the VSR1-Nt structure. Protease associated domain (light blue) and central domain (dark grey) are shown as cartoon representation. Hydrogen bonds are shown in black. The peptide is coloured in orange for CBS1 and yellow for CBS2. **A and B)** Unbiased density (green) calculated using Polder Maps at each CBS. **C)** A poly-Ala chain modelled into the 2Fo-Fc electron density (blue), showing hydrogen bonding of the C-terminus and Arg95 as well as peptide to VSR1-Nt backbone interactions. **D)** The ssVSD (Ser3-Thr10) modelled into the 2Fo-Fc electron density (blue), showing hydrogen bonding between the peptide and CBS2. **E and F)** Surface representation of VSR1-Nt coloured to represent electrostatic potential to show each modelled peptides fit within each CBS.

### VSR-Nt presents two peptide binding sites

Unbiased difference density was observed at two putative cargo-binding sites. Evidence for peptide density was strongest for cargo binding site 2 (CBS2) which is located within the central domain. Additional density at cargo binding site 1 (CBS1), located across the PA and Central domain, was consistent with peptide binding and overlapped with the cargo-binding site identified in previous PA-domain structures^10,22^ (**Fig. 3**).

Electron density consistent with CBS1 was observed within a cavity composed by switch I, the loop _95_RGDCYF_100_ and switch III in the PA domain together with _367_TI_368_ of the central domain (**Fig. 3A, C**). Although the density did not permit the unambiguous assignment of amino acid side chains, potentially reflecting conformational heterogeneity or binding in multiple orientations, a peptide backbone could be modelled into the density with favourable hydrogen-bonding interactions between VSR1 main chain atoms and the cargo peptide (ssVSD) (**Fig. 3C,E**).

CBS2 was unambiguously identified within the Central domain, around a hydrophobic groove containing the loop _234_WYCPEAFTLSKQCK_247_ (**Fig. 3B,D**). Eight residues from the cargo peptide were fitted in CBS2 and mostly engaged in hydrophobic interactions involving _234_WY_235_ and Phe240. In addition, the interaction with the peptide is stabilised by hydrogen bonds between the VSR backbone and Arg7 and Thr10 of the peptide, and Gln245 side chain and peptide backbone, and between the side chains of Gln249 and Asn4 of the NPIR motif. CBS2 is close to Ca2 and can only be formed after rearrangement of VSR upon calcium binding (**SI Fig. 3**).

### VSR1-N_t_ forming a complex with its physiological cargo in a 1:1 stoichiometry

To determine if the two cargos could be bound simultaneously or if the presence of two peptides in the structure was a consequence of using a small 10-residue peptide, we investigated the cargo:receptor stoichiometry using SEC-MALS and a recombinant Aleu-GFP construct mimicking physiological vacuolar cargo. This fusion protein contains the entire sorting sequence of the natural VSR cargo ‘aleurain’, fused to GFP, with an approximate size of the native cargo protein (∼32 kDa). Consistent with this, the addition of Aleu-GFP to VSR1-N_t_ produced a 27 kDa increase in molecular mass as measured by SEC-MALS (**Fig. 4**). This corresponds with VSR1-N_t_ forming a complex with its cargo in a 1:1 stoichiometry. This result indicates that VSR1-Nt binds a single Aleu-GFP molecule, suggesting that a single physiological ssVSD cargo molecule engages one receptor molecule at a time.

**Figure 4.**
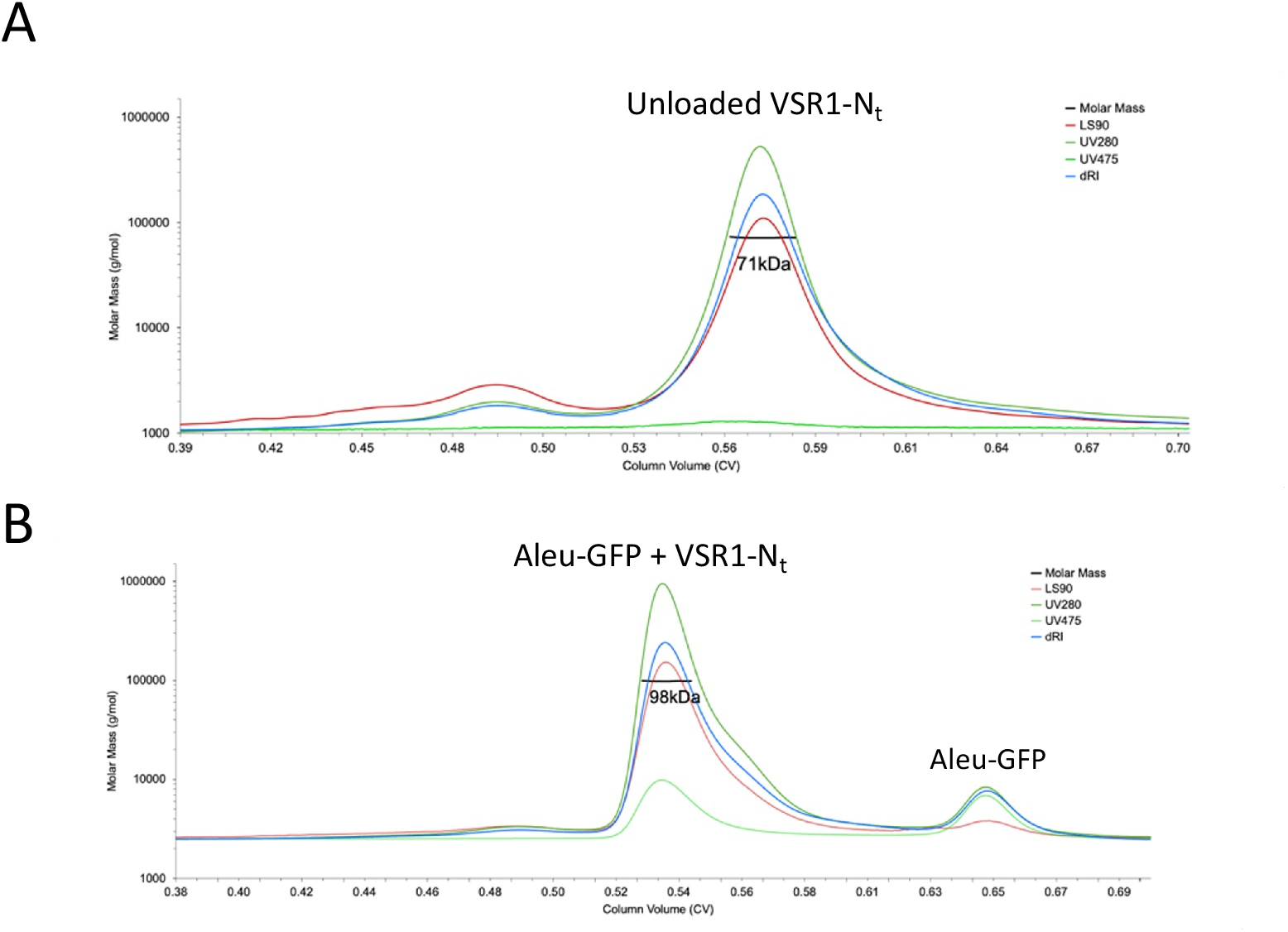
VSR1-N_t_ binds the Aleu-GFP in a 1:1 stoichiometry.. **A.** Purified unloaded VSR1-N_t_ lumenal domain was found mostly as a monomer in the presence of 1 mM calcium. **B.** Incubation of VSR1-N_t_ with cargo protein Aleu-GFP results in a size shift demonstrating a 1:1 stoichiometry receptor:cargo

### Cargo binding induces changes in solvent accessibility of specific residues

To investigate which regions of VSR1-Nt are affected by cargo binding in solution, we employed X-ray footprinting mass spectrometry (XFMS) to map *in-solution* residue-specific changes in VSR1-Nt solvent accessibility upon Aleu-GFP binding^30,31^. High-flux-density X-rays were used to generate hydroxyl radicals, which oxidatively modify solvent-exposed side chains. By varying X-ray exposure times, we generated dose-response curves for both unloaded and cargo-loaded VSR1-Nt. LC-MS/MS analysis achieved complete sequence coverage and confirmed reproducible, dose-dependent modifications. The resulting dose-response plots (unmodified fraction vs. exposure time) (**SI Fig. 4**) followed pseudo-first-order kinetics, allowing for the calculation of site-specific reactivity rate constants (*k* s^−1^. **Sl Table 5**). Because *k* is governed by both a residue’s intrinsic reactivity and its solvent accessibility, the ratio of modification rates between the free and bound states provides a ratiometric measure of accessibility changes at conformational hotspots (**Fig. 5A**).

**Figure 5.**
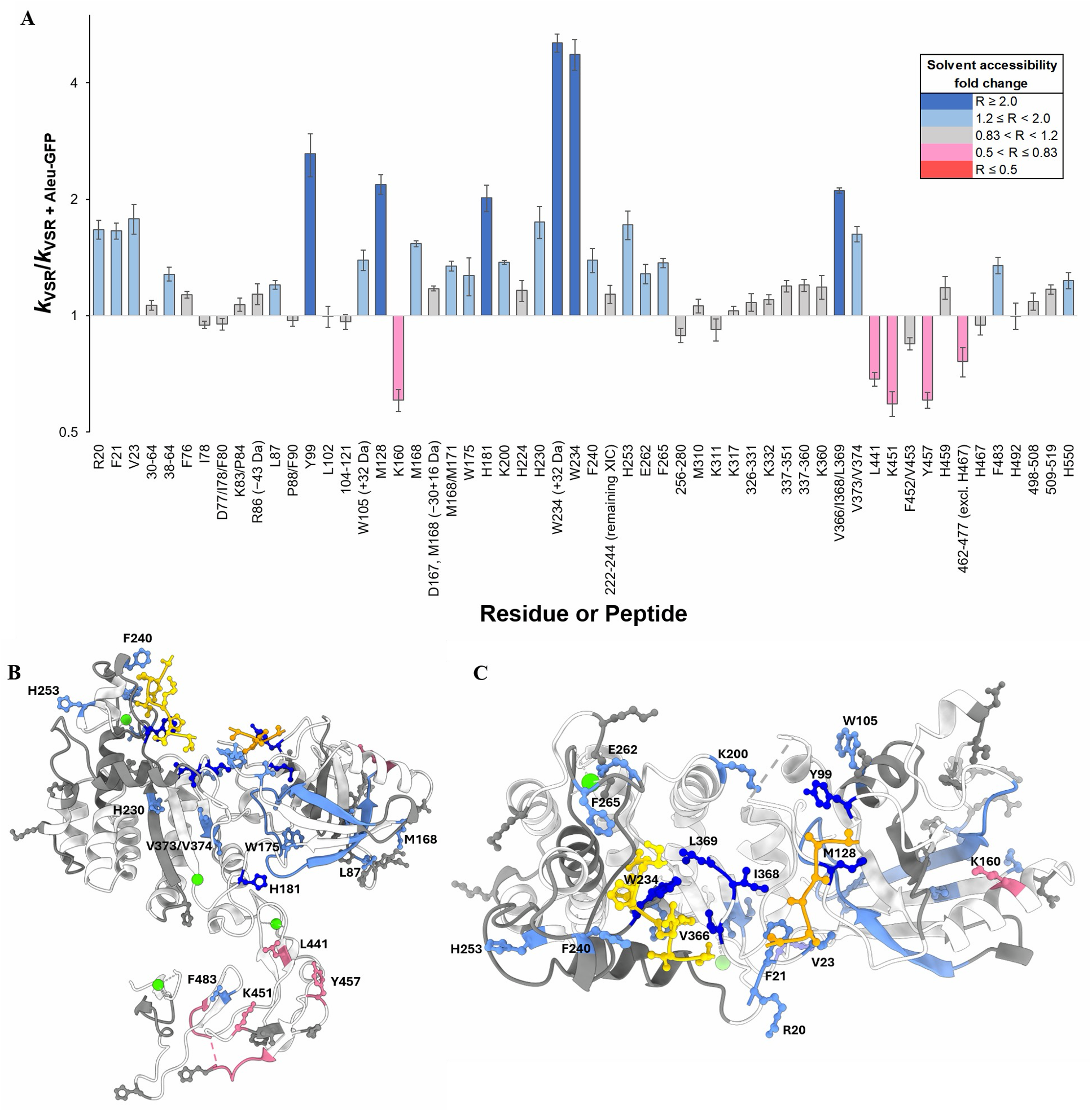
XFMS analysis of changes in VSR1-Nt solvent accessibility upon binding to Aleu-GFP. **A)** The height of each column corresponds to the ratio (R) of the hydroxyl radical reactivity rate constants of free VSR1-Nt to VSR1-Nt bound to Aleu-GFP. The site-specific ratio of rate constants measures the relative change in solvent accessibility. A ratio greater than 1 represents a decrease in solvent accessibility, while less than 1 represents an increase in solvent accessibility. Unless noted, the oxidative modification represents a hydroxylated product with a +16 Da mass shift. Co-eluting, modified peptide isomers are shown as mixed modifications or peptide-level data. Grey-colored bars indicate modification was observed but minimal change in solvent accessibility. Error bars represent the SD of the ratio. VSR1-Nt. **B)** Mapping of XFMS-derived changes in solvent accessibility onto the cargo-bound structure of VSR1-Nt. Regions in white have no associated data. The bound cargo is coloured orange (CBS1) and yellow (CBS2). Where residue-resolution data is available, side-chains are shown. **C)** Orientation from above showing CBS2 (left) and CBS1 (right).

The largest decreases (> 2-fold) in solvent accessibility upon cargo binding was observed at residues Tyr99, Met128, His181, Trp234 and Val366/Ile368/Leu369 (**Fig. 5A**). When mapped to the cargo-bound structure (**Fig. 5B and C**), Tyr99, Met128 and the Val366/Ile368/Leu369 loop belong to the CBS1 cavity, as described above (**Fig. 3A, B**). In parallel, Trp234 is part of the hydrophobic groove of CBS2 with a described hydrophobic contact (**Fig. 5B**). Moderate (2 - 1.2 fold) decreases in solvent accessibility were found at Arg20, Phe21, Val23, all belonging to Switch I of CBS1 as previously reported^22^. Moderate decrease was also found at Phe240, which belongs to the CBS2 hydrophobic groove, and Phe265 which is in close proximity to the CBS2 cargo. The decrease in rate constants at these residues are attributed to shielding by cargo binding directly to or proximal to these residues, suggesting that ssVSD cargo could potentially bind at either CBS1 or CBS2. Beyond the cargo binding sites additional residues within the PA domain (Leu87, Trp105, Met168), Switch IV (His181) and Central domain (Lys200, His230, His253, Glu262) also showed decreases in solvent accessibility, whereas the solvent accessibility of Lys160 in the PA domain is increased (**Fig. 5A**). His181 belongs to structural Switch IV involved in remodelling VSR1-Nt (**Fig. 2 C,D**). Trp105 and Lys200 are proximal to Switch III and, to a lesser extent, to residues such as Tyr99 that contribute to CBS1. Switch III is dynamic in the unloaded VSR1-N_t_ but involved in the positioning of residues such as Met128 during cargo binding, restricting Switch III dynamics, in turn impacting the dynamics and solvent accessibility of neighbouring residues. His230, His252 and Glu262 are all within 10 Å of CBS2 and experience subtle changes in their dynamics or conformation upon cargo binding, in particular Glu262 which is within Switch V. Interpretation of the moderate changes in Leu87, Met168, and Lys160, which are within a surface of the PA domain that is distal to CBS1 (**Fig. 5B**), is harder to interpret and may reflect a more complex conformational change.

Additional changes were observed within the EGF domains. Residues Leu441, Lys451, Tyr457 and the 462-477 region (excluding His467) of EGF1 experience a moderate increase in solvent accessibility, while Phe483 and His550 of EGF2 and EGF3 respectively experience a moderate decrease in solvent accessibility (**Fig. 5A and B**). The increase in solvent accessibility in EGF1 is consistent with its observed movement away from the central domain in the cargo-bound state (**Fig. 2C**). Since these residues are far from CBS1 and CBS2, the changes in accessibility reflect induced structural changes translated to the EGF domains upon cargo binding. This is compatible with various hypotheses such as that EGF domains have a role in modulating cargo binding, or alternatively these observed conformational changes are the result of allosteric communication involved in signal transduction from cargo binding in the PA and Central domain to the EGF domains^15^.

Together the SEC-MALS and XFMS analyses confirm that VSR-Nt binds a single physiological cargo molecule, while undergoing large structural rearrangements. Cargo binding not only affects the accessibility of residues in both CBS1 and CBS2 but also influences accessibility of residues in the EGF domains, suggesting that EGF domains also participate in structural changes upon cargo binding.

### Calcium and pH control VSR1-Nt conformation and cargo binding *in solution*

To better understand how changes of lumenal pH and calcium concentrations encountered during VSR1-N_t_ trafficking influence the receptor conformation and cargo binding, we performed SEC-SAXS analysis under conditions mimicking early (neutral pH, high calcium) and late (acidic pH, low calcium)) compartments^32,33^. Since the organelle calcium concentrations are not well established, we opted for pH 7.4 with 1 mM calcium for early compartments and pH 5.6 with no calcium for late compartments conditions. We also analysed specifically the effect of the presence, and absence of calcium on the structure of VSR1-N_t_ at fixed pH (pH 5.6 with no calcium vs 5 mM calcium) (**Fig. 6**).

**Figure 6.**
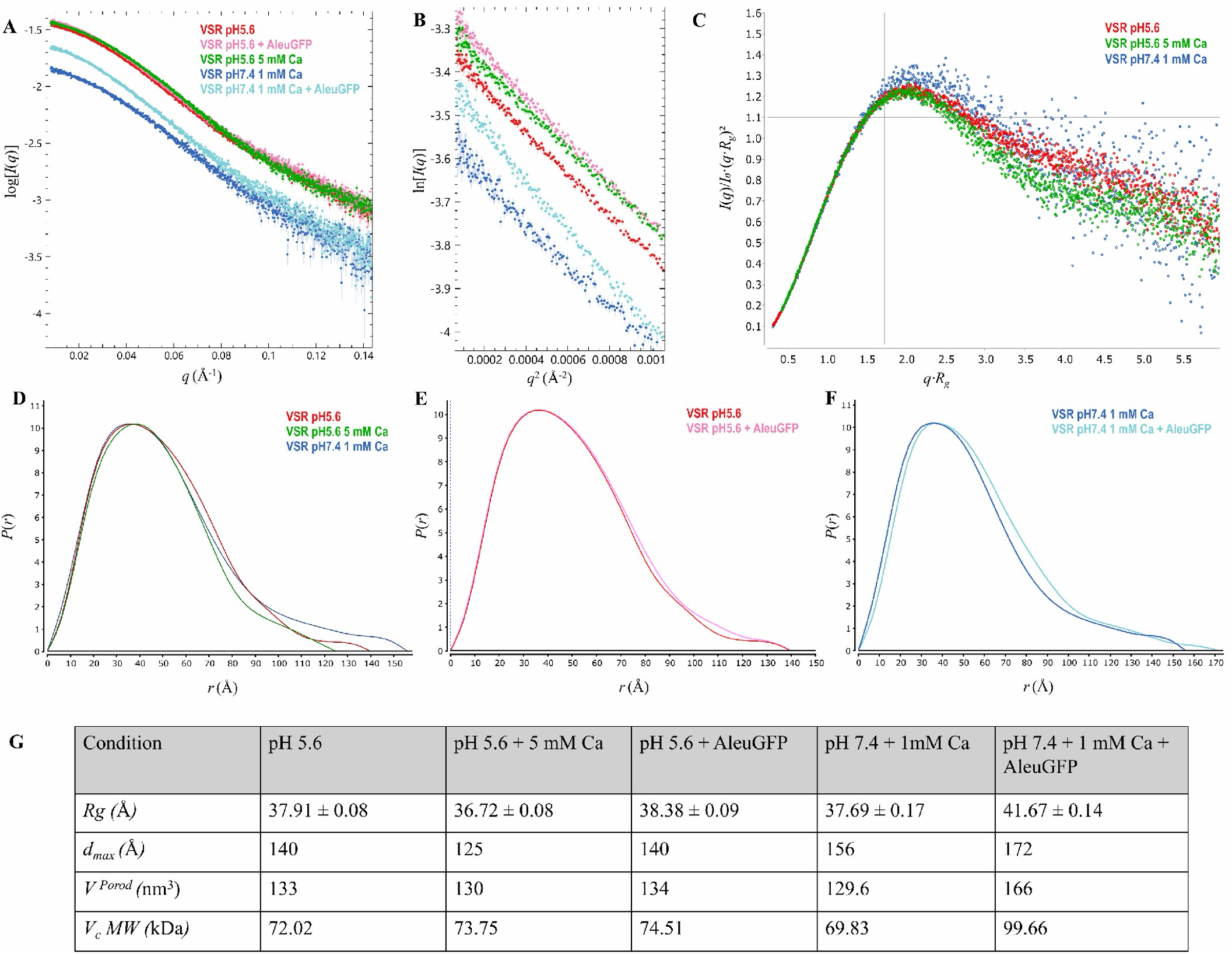
SAXS analysis of VSR-Nt in different pH and calcium conditions and in the presence or absence of Aleu-GFP Cargo. **A)** Log10 SAXS scattering profiles for each condition. **B)** Guinier region, from which *Rg* is estimated. **C)** Dimensionless Kratky plot, the intersection of the black trace indicates the theoretical peak position of a globular protein. **D-F**) Pair-distance distribution *P(r)* function, used to determine particle shape and *Dmax*. **G)** Table summary of SAXS derived structural parameters. V_Porod_ is Porod Volume, and molecular weight (MW) derived from Volume of Correlation.

At conditions mimicking late compartments (acidic pH, no calcium), VSR1-N_t_ forms a single monodisperse species in solution (**SI Fig. 5A**), with radius of gyration (*Rg*) of 37.91 ± 0.08 (**Fig. 6B, G**) and maximum dimension (*Dmax*) of 140 Å (**Fig. 6D, G**). Kratky analysis (**Fig. 6C**) and Porod exponent of 3.1 indicate flexibility, confirmed by the extended tail on the pair-distance distribution *P(r)* function (**Fig. 6D**), likely representing the EGF repeats attached to a more restricted Central and PA domain. The addition of calcium induces a reduction of *Rg* to 36.72 ± 0.08 Å (**Fig. 6B, G**), *Dmax* to 125 Å, a reduction in flexibility evidenced by the Kratky plot (**Fig. 6C**) and compaction in the *P(r)* distribution, consistent with a more compact conformation and EGF tail (**Fig. 6D**). At conditions mimicking early compartments, (neutral pH with calcium), there was an increase in *Rg* to 37.69 ± 0.17 Å (**Fig. 6B, G**) and *Dmax* to 156 Å, explained in the *P(r)* distribution by the extension of the flexible EGF tail, while the core remains in the calcium-dependent compact conformation (**Fig. 6D**).

When Aleu-GFP cargo is added to VSR at late compartment conditions (i.e acidic pH), a shoulder peak of a species smaller than VSR1-N_t_ is observed by SEC-SAXS (**SI Fig. 6C**) consistent with free Aleu-GFP peak, and no significant changes observed in the VSR1-N_t_ peak (*Rg* 38.38 ± 0.09 Å; *Dmax* 140 Å; similar *P(r)* distribution) (**Fig. 6B, E, G**) indicating no interaction at this condition, as expected in the absence of calcium. In contrast, at early compartment conditions ( i.e pH 7.4 and 1mM calcium), a shift in elution time of the VSR1-N_t_ peak is observed (**SI Fig. 6F**), and increases in *Rg* to 41.67 ± 0.14 Å, *Dmax* to 172 Å, Porod volume to 166 nm^3^, Volume of Correlation *V_c_* derived molecular weight increase of 30 kDa consistent with the SEC-MALS data, and positive shift in the *P(r)* distribution (**Fig. 6B, F, G**). Altogether, cargo binding is confirmed in this condition, in agreement with previous results that the presence of calcium ions are essential for cargo binding and neutral pH favours this interaction.

To further model in-solution shape and domain organisation, *ab initio* molecular envelopes of VSR1-N_t_ at both late (pH 5.6) and early (pH 7.4 + 1mM Ca) conditions were computed from the SAXS data (**Fig. 7**)^34^. The VSR1^pH5.6^ envelope (χ^2^=1.16) adopts an elongated structure with a core spanning 109 Å (**Fig. 7A**). In comparison, the envelope of VSR1^pH7.4, 1mM^ ^Ca^ (χ^2^=2.546) contains a core spanning 97 Å with a cleft between two lobes, and a more narrow and elongated tail (**Fig. 7B**). CIFSUP alignment of SASREF structural models (pH 5.6 χ^2^=3.06; pH 7.4, 1mM Ca χ^2^=3.964) to the molecular envelopes indicates the extended tails are occupied by the EGF domains, with cores composed of PA and Central domains (**Fig. 7C-D**). The VSR1^pH7.4, 1mM^ ^Ca^ SASREF model contains a cleft between the PA and Central domain, whereas the VSR1^pH5.6^ SASREF model positions these domains further apart (**Fig. 7E**). The domain spacing of the VSR1^pH^ ^5.6^ SASREF model closely resembles the unloaded VSR1-N_t_ crystal structure (**Fig. 7F**). The domain spacing in the VSR1^pH7.4, 1mM^ ^Ca^ SASREF model resembles the calcium bound, cargo-loaded VSR1-N_t_ crystal structure (**Fig. 7G**).

**Figure 7.**
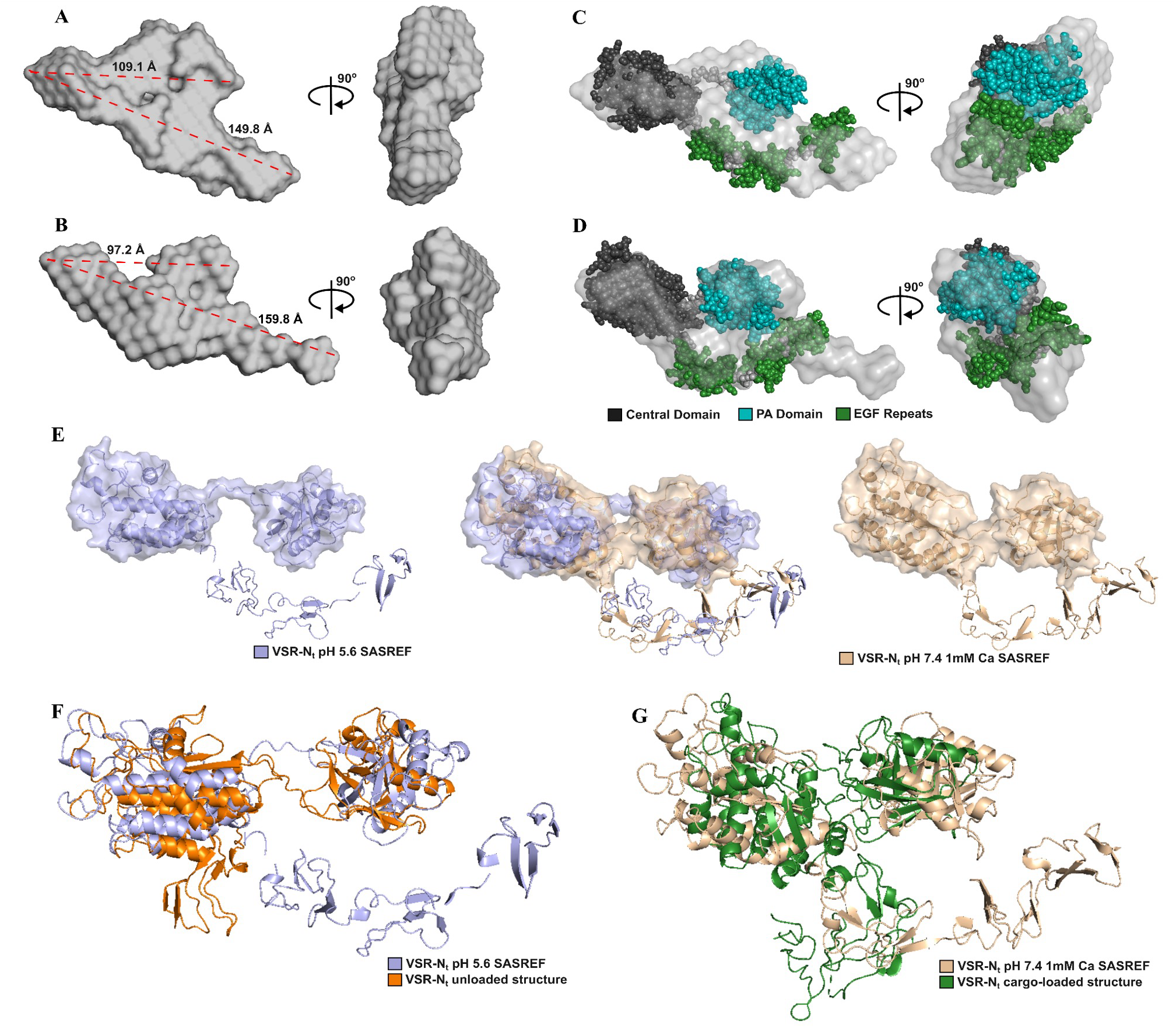
Superimposition of SAXS envelopes with ab initio models. **A-B**) DAMMIF/DAMAVER/DAMMIN-derived molecular envelopes of VSR-Nt at pH5.6 and pH7.4, 1mM Ca respectively, aligned to two *Dmax* vectors of the core and the entire envelope. **C-D**) Fitting of *ab initio* SASREF structural model to molecular envelopes, for VSR at pH5.6 and pH 7.4, 1mM Ca respectively. Models are orientated by alignment of SASREF structures. **E)** Superposition of aligned SASREF structural models. Surfaces are shown for the Central domain, PA domain and connecting Switch IV. **F)** Superimposition of the pH5.6 SASREF model and the unloaded crystal structure. **G)** Superimposition of the pH7.4, 1mM Ca SASREF model and the calcium bound cargo loaded crystal structure.

Together these data suggest that the decrease in lumenal pH and calcium concentration encountered during VSR1 trafficking promotes a conformational transition from a compact, cargo-binding competent state towards a more open conformation resembling the unloaded structure. This transition is consistent with the established role of acidic, low-calcium compartments in cargo release.

## Discussion

### A calcium-dependant structural mechanism for cargo recognition and release by VSR1-Nt

The mechanisms by which calcium and pH regulate cargo recognition and release during VSR trafficking have remained longstanding questions in plant cell biology^15^. Comparison of the unloaded and cargo-loaded VSR1-Nt crystal structures presented here provide a structural explanation.

The cargo-bound structures revealed four calcium-binding sites distributed throughout the lumenal domain, with two sites located in the Central domain and one present in each of the two resolved EGF repeats. In contrast, no biologically relevant calcium-binding sites were identified in the unloaded structure. Calcium coordination is accompanied by large-scale rearrangements involving seven conformational switches spanning the PA, Central and EGF domains. These include the previously described PA-domain switches I-IV^10^ and three newly identified switches (V-VII). Importantly, calcium dependent remodelling of switches I, III, IV, V and VI results in the reorganisation of CBS1 and exposure of CBS2 (**SI Fig.3**). Thus, calcium binding could act as a structural trigger that primes VSR1 for cargo binding.

The relevance of the structural observations from the unloaded and loaded structures were further supported and complemented by *in solution* studies. SEC-SAXS analysis demonstrates that calcium alone induces the more compact state bringing the PA and Central domains into close proximity while simultaneously rigidifying the EGF region into a more restricted and less dynamic state. The severity of EGF restriction shows sensitivity to changes in pH, implying a hypothetical role as a pH sensor. Interestingly, Glu414 which is involved in a salt bridge in the unloaded structure between the CD and EGF domain, is engaged in calcium binding of Ca3 in the cargo-bound form. Cargo binding was only observed under calcium-containing conditions, confirming that calcium-dependent remodelling is required for cargo recognition. Together, these findings suggest that calcium does not merely enhance ssVSD cargo affinity but directly controls the structural transition between ssVSD cargo-binding incompetent and competent receptor states. While this could also be compatible with ctVSDs, further investigation is required to confirm whether this is true.

The impact of calcium and pH on VSR cargo binding cycle is reminiscent of a mode of action called the pH relay hypothesis. In this model, changes in pH modulate calcium affinity and thereby influences receptor conformation and cargo binding^35^. Under high calcium conditions, changes in pH are expected to have relatively little effect because calcium saturation stabilises the binding sites. However, under low-calcium conditions, protonation of residues involved in calcium coordination would favour calcium release and consequently destabilise the cargo-binding conformation. For VSR1, such conditions are compatible with cargo release in late endosomal compartments where both calcium concentration and pH decrease^36,37^. In agreement with this, previous purifications of pea VSR using a cargo-bound column, performed in the presence of calcium at neutral pH, required a mildly acidic conditions (pH 4) and the presence of EGTA for VSR to be released from the column, supporting the proton relay hypothesis as a control mechanism for cargo affinity^37^.

The C-type lectin receptor langerin has been shown to rely on a proton relay mechanism involving a specific histidine residue (His294) located in a loop nearby to the cargo binding domain^38^. Interestingly, three histidine residues (Histidine 181 and Histidine 230 and 253) undergo a mild decrease in accessibility when transitioned from unloaded-to the cargo-loaded form (**Fig. 5**). His181 is located in switch IV between the PA and the Central domain, close to Ca1 and forms a salt bridge with Glu177 in the cargo loaded form presented here. Switch IV is involved in the large-scale calcium dependent rearrangements and shows a high degree of sensitivity to both cargo-induced and calcium-induced allosteric communication. Further experimental work would be required to decipher if specific residues in VSR1-Nt, such as these histidine residues, could be sensitive to pH changes modulating calcium binding and VSR structure rearrangement.

### Discovery of a novel cargo-binding site in the central domain

Previous structural studies established the PA domain as the principal cargo-recognition region of VSR1 and identified CBS1 as the primary binding pocket for vacuolar sorting determinants^10,11,22^. The structures presented here expand this model by revealing a second cargo-binding site (CBS2) within the Central domain. Although a role for the Central domain in cargo recognition has previously been proposed^8^, the well-defined peptide electron density observed at CBS2, together with XFMS analysis using the cargo Aleu-GFP, supports the conclusion that CBS2 represents a true cargo-binding site rather than a crystallographic artefact.

CBS2 is positioned adjacent to the Ca2 calcium-binding site and is formed through a calcium-dependent rearrangement of newly identified Switch V. In the unloaded structure, this region participates in intermolecular contacts within the trimeric assembly. Upon calcium binding, however, it undergoes a substantial conformational change that exposes CBS2 as a well-defined peptide-binding pocket. These observations indicate that calcium binding not only drives global structural rearrangements in VSR but also directly exposes and stabilises a previously inaccessible cargo-binding surface within the Central domain.

The identification of CBS2 refines the model proposed by Luo *et al*.¹⁰, who suggested that calcium-induced conformational changes within the PA domain and Switch IV reposition the Central domain to face the CBS1 site and facilitate recognition of the NPIR-containing sorting determinant. While CBS1 was proposed to contribute to ssVSD and ctVSD binding in a crystal structure containing only the PA domain^10,11,22^, our structures identify CBS2, in the central domain, as the preferred binding site for the ssVSD peptide. Notably, unambiguous electron density is observed for the broader SNPIRPVT sequence rather than for the NPIR motif alone, indicating that residues flanking the NPIR motif also contribute to cargo recognition. This finding suggests a more extensive recognition interface than previously appreciated. Consistent with this interpretation, residues surrounding CBS2 exhibit reduced solvent accessibility following addition of Aleu-GFP further supporting a direct role for CBS2 in cargo engagement.

Although CBS2 displays the most well-defined ssVSD-binding density, we also observe electron density for peptide at CBS1 in a position similar to that previously reported^10^. However, unlike CBS2, the density at CBS1 could not be unambiguously assigned to the ssVSD peptide. Consequently, a poly-alanine model was used to represent the observed density (**Fig. 3C**). Despite this limitation, the density is suggestive of a binding mode similar to that described previously, including a potential interaction between Arg95 and a C-terminal carboxyl group.

The weak and poorly defined nature of the density suggests that NPIR-containing peptide binding at CBS1 may be promiscuous. Such promiscuity may explain why ssVSD-containing cargo can occupy this site, while also indicating that CBS1 is unlikely to represent the ssVSD recognition site. Previous structural studies modelled ctVSD peptides within CBS1, and the best-fitting orientation of the density observed here is likewise consistent with direct engagement of a free C-terminal carboxyl group by Arg95. Such an interaction would not be possible for ssVSD containing cargo, as ssVSD signal sequences are located at the N-terminus of cargo proteins and therefore lack a free C terminus as part of the signal sequence. Therefore, our observations support a more specialised role for CBS1 in the recognition of C-terminal vacuolar sorting determinants (ctVSDs) instead. This interpretation is further supported by the fact that the plant Receptor Homology– Transmembrane RING-H2 (RMR) proteins, which contain a PA domain including CBS1 but lack the Central domain, have been implicated in the transport of ctVSD-containing cargos but not ssVSD^39^. Taken together, these observations suggest a functional division between the two binding sites, with CBS1 preferentially recognising ctVSD-containing cargoes and CBS2 serving as the primary recognition site for ssVSD-containing cargoes.

### Functional implications of the two cargo-binding sites

The discovery of CBS2 suggests that cargo recognition by VSR1 involves a more extensive binding interface than previously appreciated. Rather than functioning exclusively through the PA-domain binding pocket, VSRs may employ coordinated interactions between the PA and Central domains. Supporting this hypothesis, XFMS analysis using Aleu-GFP detected protection patterns around both CBS1 and CBS2, indicating that both regions participate in cargo engagement in solution.

At first glance, the observation of two bound peptides in the crystal structures appears inconsistent with SEC-MALS experiments showing formation of a 1:1 VSR1:cargo complex. However, these findings can be reconciled by considering the nature of the ligands used. While the short synthetic peptides employed for crystallisation are sufficiently small to occupy both sites simultaneously, physiological cargo proteins present sorting determinants within the context of larger folded domains and steric constraints may therefore limit concomitant occupancy. This is also suggested by the SEC-MALS experiments showing that VSR binds to only one Aleu-GFP. Alternatively, interaction with one site may allosterically influence accessibility of the second. In fact, several non-mutually exclusive functional models can be envisaged. First, CBS1 and CBS2 may display preferential specificity toward different classes of vacuolar sorting determinants. Alternatively, the two sites may exhibit different intrinsic affinities, enabling recognition of the highly diverse sorting determinants found in vacuolar cargo proteins. Third, CBS1 and CBS2 could act sequentially or cooperatively during cargo capture, with initial binding at one site promoting stabilisation of interactions at the second.

Current data do not distinguish between these possibilities. However, the combined crystallographic, SAXS, SEC-MALS and XFMS observations strongly support the conclusion that both CBS1 and CBS2 contribute to cargo recognition. Quantitative binding studies and *in planta* studies using native ssVSD- and ctVSD-containing cargoes will be required to define their precise functional roles.

### Role of EGF domains in structure stabilisation and signal transduction

While calcium binding induces a rearrangement that exposes cargo binding sites in the PA and Central domain, our findings suggest that EGF repeats play a role in structural stabilisation of the calcium bound structure and that the structural effects of cargo binding further extends throughout the lumenal region of VSR1-Nt, including the EGF domains.

Calcium binding sites were identified within both resolved EGF1 and EGF2 repeats, and calcium binding alters not only the relative positioning of the PA and Central domain, but also the overall conformation and flexibility of the EGF repeats, as seen by SEC-SAXS analysis. These observations suggest the EGF repeats contribute to sensing and responding to changes in calcium. In this context, the EGF domains may act as structural modulators that stabilise cargo-bound conformations under calcium-rich conditions. This interpretation is consistent with previous findings that calcium binding by EGF repeats stabilise receptor-cargo interactions^15^.

In addition, XFMS derived changes in solvent accessibility are detected within the EGF repeats, coupling cargo recognition with long-range conformational rearrangements. This would be compatible with a role of the EGF domains in signal transduction, from cargo binding in the PA and Central domains to the membrane-proximal and transmembrane region^40^.

The EGF conformational plasticity may therefore reflect a role in coupling cargo binding and VSR trafficking. In this context, VSR has been suggested to exist in multiple forms in cells, including a high molecular form^41^. Homomerisation was suggested to be driven by both the transmembrane and C-terminal domain and is crucial for VSR function *in vivo* ^20,41^. Despite the lack of these domains in their constructs, Park *et al.,* have recently reported the X-ray structure of an unloaded form of VSR1-N_t_ that crystalises as a trimer. The authors propose a functional role for this trimer, in line with *in vivo* observations^21,41^. Consistent with these observations, we solved an X-ray structure of trimeric unloaded VSR1-Nt (**Fig. 1C**) while our cargo loaded structures was solved as a monomers. This may reflect cargo-induced or calcium-induced changes in physiological VSR1 inserted in the membrane. Since our attempts to purify full length VSR1 including the transmembrane domain and C-terminal tail were not successful, further *in planta* experimental work based on our structural results will allow validation of this hypothesis.

### Overall conclusions

Taken together, the crystallographic, XFMS, SEC-MALS and SAXS data support a model in which VSR1 functions as a highly dynamic and integrated receptor, with the PA, Central and EGF domains acting cooperatively rather than as independent modules. Calcium binding drives extensive receptor remodelling, generating a cargo-binding competent state and exposing a previously unresolved cargo-binding site within the Central domain. Our findings identify two spatially distinct cargo-binding sites, with CBS2 emerging as the principal recognition site for ssVSD-containing cargoes, while CBS1 may play a specialised role in the recognition of ctVSD-containing cargoes. Cargo binding is accompanied by long-range conformational changes extending into the EGF domains, suggesting that these domains contribute to stabilisation of the calcium-bound state and may participate in allosteric communication across the receptor. As VSR1 traffics into increasingly acidic and calcium-depleted late compartments, the receptor transitions towards a more open conformation with reduced cargo affinity, promoting cargo release. Together, these findings provide a structural framework for understanding how lumenal calcium and pH regulate cargo recognition, trafficking and release during plant vacuolar protein sorting.

## Materials and Methods

The methods are described briefly here but more detail can be found in SI material and methods.

### Constructs

VSR1 (AT3G52850) was amplified from *Arabidopsis thaliana* genomic DNA and cloned in a pUC vector. The lumenal region of the *VSR1* gene (VSR1-N_t_) was further amplified and inserted into the pOPIN-HBM vector (Oxford Protein Production Facility, OPPF) flanked by HBM signal, and a C-terminal His_8_ tag. Aleurain (Aleu) construct was amplified and cloned as a GFP fusion protein in a pUC vector to generate the pJA1 vector. The Aleu-GFP cassette was then subcloned into the pET28b (Novagen). All constructs were sequenced at each stage of cloning to confirm identity. A list of primers is presented in **SI Table 3**.

### Expression and purification of VSR1-N_t_ and Aleu-GFP

The lumenal N-terminal region of VSR1-N_t_ was expressed in Expi293 cells in the presence of kifunensine 1 mg/ml following the published protocol^42^. Thirty ml of Expi cell culture resulted in a yield of 4.25 mg of VSR1. This was concentrated to 15 mg/ml and used for further crystallization studies. For SEC-SAXS, sample “VSR1-N_t_ pH 7.4, 1mM Ca”, purification was as above but the final size-exclusion chromatography (SEC) purification was into 20 mM HEPES pH 7.4, 200 mM NaCl, 1 mM CaCl_2_ and 1 mM MgCl_2_. For sample “VSR1-N_t_ pH 5.6”, the IMAC buffer was 20 mM Tris pH 7.4, 200 mM NaCl. The IMAC elution fraction was buffer exchanged by PD-10 into 20 mM MES pH 5.6, 200 mM NaCl and purified by SEC on Superdex 200 10/300. Calcium was then directly added for the “VSR1-Nt pH 5.6, 5mM Ca” sample.

Aleu-GFP was expressed in *E.coli* BL21 Star (DE3) (ThermoFisher) in autoinduction media at 37° C for 16 hours^43^. The cells were lysed and the protein was purified through affinity purification with Ni-NTA agarose beads. The Aleu-GFP was further purified with size exclusion chromatography via ÄKTA pure protein purification system (Cytiva) using Superdex 200 Increase 10/300 GL column (Cytiva) using 1.5 CV in 20 mM Tris/HCl pH 8.0, 200 mM NaCl (with or without 1mM CaCl_2_ and 1 mM MgCl_2_). Fractions containing the fluorescent protein were pooled and the protein was concentrated with Vivaspin 20 (10 kDa MWCO) to the desired concentration.

### Crystallographic determination of unloaded VSR1-N_t_

Purified VSR1-N_t_ was set up for crystallization trials at 15 mg/ml initial concentration with a 1:1 protein to well solution ratio and a total drop volume of 200 nl. VSR1 crystals grew after 13 days at 4° C with reservoir solution 0.1 M Tris/HCl pH 8.0, 20% v/v glycerol ethoxylate, 10% v/v tetrahydrofuran and frozen with no cryoprotectant in liquid nitrogen. Data collection was performed at beamline i03 at the Diamond Light Source. Diffraction data was processed with XDS^44^ to 3.51 Å resolution and P2_1_3 space group with cell constants: a = b = c = 141.3 Å.

A partial crystallographic solution of VSR1 crystallographic was obtained by molecular replacement in PHASER^45^ using the complex state of the PA domain (PDB id: 4TJX)^10^ as the search model. The Central domain was phased using a low-resolution methodology within the ARCIMBOLDO principle: expansion of accurate partial structures through density modification and map interpretation with multiple fragment hypotheses evaluation with and without side chains^46,47^. Multiple fragments were placed in real space with Coot^48^ using the density modification map calculated with SHELXE^49^ from the partial solution containing the PA domain. Alternative structural hypotheses were evaluated considering different secondary structure elements, fragment reversion and side chain atoms inclusion using the most probable sequences generated with the SEQUENCE SLIDER functions^50,51^. Fragments were scored with PHASER log-likelihood gain^45^. A model of VSR1 predicted by AlphaFold^52^ using GoogleColab and from the joint database with EBI was used to confirm the Central and EGF domain modelling. The model was built using manual building in Coot^53^ and refinement with PHENIX^54^. Each glycosylation was supported by stereochemical and real-space correlation coefficient validation, in agreement with expected values calculated with Privateer^26^.

To evaluate potential presence of calcium ions, above and below the calcium K-edge (λ = 3.06 Å), at 3.02 Å and 3.2 Å respectively, were collected on beamline I23 at Diamond. The diffraction images were processed using the xia2 DIALS pipeline^55,56^ and the relevant sweeps were combined using xia2.multiplex^57^. Molecular-replacement solutions were generated using DIMPLE pipeline and anomalous difference Fourier maps and peak heights were subsequently calculated using ANODE^58^

### Crystallisation and structural determination of VSR1-N_t_ and peptide complex

VSR1-N_t_ was concentrated to 8 mg/ml and incubated with cargo peptide (ADSNPIRPVT synthesized by peptide synthetics) in 50 mM HEPES (7.5), 150 mM NaCl and 2 mM CaCl_2_ at 1:5 molar ratio for 30 minutes before crystallisation using sitting drop vapour diffusion at 4 °C (drop size 150 nL) using a Mosquito. The protein complex crystallised in two conditions in MemGold (Molecular Dimensions) G3 (0.1 M Tris (8.5), 0.1 M MgCl_2_, 0.1 M NaCl, 33 % (v/v) PEG 400) and E12 (0.05 M Glycine (9.5), 100 mM NaCl, 33 % (v/v) PEG 300), with a drop ratio of 1:2, protein to reservoir solution.

Crystals were harvested directly from the drop using micro mount loops, then cooled using liquid nitrogen. All diffraction data were collected at Diamond Light Source synchrotron (Diamond Light Source, UKLS), UK.

Native X-ray data were collected at 100 K at I04 microfocus beamline of the G3 crystals. Diffraction patterns were integrated using xia2 pipeline running DIALS^55,56^. The phases were determined by molecular replacement using Phenix Phaser-MR with the predicted AlphaFold3^52,59^ model trimmed (residues 20-407) as the search model. The initial electron density maps were inspected, and the model was built using manual building in Coot^53^. The structure was refined using PHENIX refine, REFMACAT and Refmac5 ^54,55,60^.

To determine whether calcium ions were present in the crystal structure, X-ray diffraction data were collected at the long-wavelength I23 beamline^61,62^, above and below the calcium K-edge at 3.07 Å and 3.09 Å respectively, on the crystals grown in E12 crystallization condition. All data were processed using xia2 DIALS, and the sweeps were combined using xia2 multiplex^57^. Anomalous difference Fourier maps and anomalous peak heights were calculated with ANODE^58^ using the molecular replacement solution generated by the DIMPLE pipeline. The anomalous peaks for each wavelength are shown in **SI Table 4**.

### SEC-MALS

SEC-MALS experiments were performed at ambient temperature, using an HPLC+LS+RI+QELS configuration pre-equilibrated with SEC buffer (20mM Tris/HCl, 200mM NaCl, 1mM CaCl_2_, 1mM MgCl_2_ adjusted to pH 7.5 at room temperature). The instrument set up included the HPLC modules of Agilent 1260 Infinity II series (Agilent Technologies) connected in-line to the miniDAWN (NEON) with a three-angle (49°, 90° and 131°) light scattering detector, and an integrated QELS dynamic light scattering detector at 135° for determination of hydrodynamic radius. Concentration was measured using UV absorbance at 280nm using the in-line 1260 infinity II series DAD detector (Agilent Technologies) and the Wyatt Optilab™ (NEON) refractive Index detector (Wyatt Technology). Sample submission, data collection and SEC-MALS analysis was performed using the HPLC connect and ASTRA 8.1.2 software (Wyatt Technology). Approximately 80µg of protein was injected onto Superdex 200 Increase 10/300 GL (GE, Life Science) SEC column at a flow rate of 0.4ml/min for run time of 70 minutes per sample. The extinction coefficients for Aleu-GFP Cargo, VSR1 and VSR1: Aleu-GFP Cargo complex was defined as 0.683 mL/mg.cm, 1.410 mL/mg.cm, and 1.165 mL/mg.cm respectively. The DAD module G7115A (Agilent Technologies) of HPLC was set to collect UV absorbance signals at 280nm, 488nm, 509nm, 395nm, 475nm to track the GFP tagged cargo. Data collection for QELS was set to DLS interval of 2 sec and Collection Interval of 0.5sec.

### Small Angle X-Ray Scattering

SEC-SAXS data was collected on Diamond beamline B21. Aleu-GFP was dialysed overnight into buffers matching VSR1-Nt, added to a 1:1 stoichiometric ratio to VSR and incubated overnight on ice. Final concentrations were 3.98 mg/mL VSR1-N_t_ in 20 mM MES pH 5.6, 200 mM NaCl; 3.92 mg/mL VSR1-N_t_ in 20 mM MES pH 5.6, 200 mM NaCl, 5 mM CaCl_2_, 5 mM MgCl_2_; 2.56 mg/mL VSR1-N_t_ with 1.29 mg/mL Aleu-GFP in 20 mM MES pH 5.6, 200 mM NaCl; 2.9 mg/mL VSR-N_t_ in 20 mM TrisHCl pH 7.4, 200 mM NaCl, 1 mM CaCl_2_, 1mM MgCl_2_; 2 mg/mL VSR1-N_t_ with 0.9 mg/mL Aleu-GFP in 20 mM TrisHCl pH 7.4, 200 mM NaCl, 1 mM CaCl_2_, 1mM MgCl_2_. Samples were injected onto a 15℃ pre-equilibrated Sepax SRT® SEC-300 column using an in-line Agilent HPLC system and data collected continuously on an EigerX 4M (Dectris) detector with 3-s exposures. The *q* range (*q*=4π*sin(θ)/λ) was 0.0045-0.34 Å^-1^, exposure temperature 288 K, and wavelength 0.9464 Å.

Buffer subtraction and merging of frames was performed using ScÅtter IV. ScÅtter IV and ATSAS 3.2.1 were used for standard data processing. *Rg,* molecular weight derived from *V_c_*, and Porod volume were determined using Primus. *P(r)* plot, *Dmax* determination, normalized Kratky plots and determination of Porod exponent were generated using ScÅtter IV. The molecular envelopes were generated using 11 iterations of DAMMIF, averaging using DAMAVER, and final refinement of the damstart output using DAMMIN. For structural modelling, the Alphafold model (AF-P93026-F1-v6) was separated into 9 regions (PA [21-173], linker 1 [174-184], CD [185-406], linker 2 [407-412], EGFR1 [413-462], linker 3 [463-467], EGFR2 [468-510], linker 4 [511-515], EGFR3 [516-554]), and modelled to the SAXS data using SASREF simulated annealing, with contact restraints to maintain domain connectivity. CIFSUP was used for model alignment to envelopes, and CRYSOL used to determine χ^2^ of envelope and model fits to experimental data. DAMMIF/DAMAVER/DAMMIN, SASREF, CIFSUP and CRYSOL are all programmes included in the ATSAS suite^63^. Data and models are available at SASBDB.

### X-Ray Footprinting Mass Spectrometry

Protein samples of VSR1-Nt alone (6 μM) and in complex with Aleu-GFP (5.5 μM VSR1-Nt, 11 μM Aleu-GFP) were dialyzed overnight at 4 °C into 10 mM ammonium bicarbonate (pH 7.7), 1 mM CaCl₂, and 200 mM NaCl. Samples were irradiated at beamline 3.3.1 of the Advanced Light Source (Lawrence Berkeley National Laboratory), which provides a high-flux 3–12 keV broadband X-ray beam as reported previously^64^. Hydroxyl radical dose variability was assessed using automated Alexa Fluor 488 fluorescence dosimetry^65^. Using the microfluidic sample delivery system, the samples were exposed for 0 – 1200 µs, quenched immediately with 10 mM L-methionine amide, and stored at −80 °C.

Exposed samples were denatured, reduced, and digested with trypsin/Lys-C (1:20, w/w, 12 h, 37 °C). The digested samples were desalted using a C18 spin column under acidic and reducing conditions, dried, and reconstituted in 0.1% formic acid. Reverse phase and high-flow bottom-up LC–MS/MS was performed on a Vanquish Flex UHPLC coupled to an Orbitrap Exploris 480 (Supporting Method).

Peptides were identified and quantified using Byos v5.10 (Protein Metrics) with 10 ppm precursor (MS1 spectra) and 20 ppm fragment tolerances (MS2 spectra), allowing up to two missed cleavages and variable oxidative modifications (+14, +16, +32, and +48 Da). The Byos-assigned site-specific modifications were validated manually from MS2 spectra. Based on extracted ion chromatograms, the software automatically calculated the fraction unmodified for each peptide as the ratio of the unmodified peptide peak area to the peak areas of the sum of unmodified and modified peptides. The dose-response data representing the fraction unmodified as a function of exposure time were fitted to a first-order exponential function, y = e−kt, to determine the hydroxyl radical reactivity rate constant, k (s−1). We reported rate constants only for fits with R² values between 0.90 and 0.99 to exclude contributions from secondary reactions (Supporting Method).

### Inclusion and Ethics Statement

All authors included in this study have fulfilled the criteria for authorship required by Nature Portfolio journals. All authors are committed to promoting equality, diversity, and inclusion, as well as fostering environments where all individuals, regardless of background, identity, or perspective, are respected and valued. This paper reflects our dedication to creating a scientific community that ensures equitable opportunities for collaboration and embraces diversity and inclusivity in every aspect of our research endeavours.

## Code availability statement

The authors are committed to share the code and mathematical algorithm used in this study upon email request to the corresponding authors. Programs named are distributed through CCP4, so accessibility is granted.

## Data availability statement

The atomic coordinates have been deposited in the Protein Data Bank (PDB identifier [ID] code: 8R4Y unloaded and 34RZ cargo-loaded).

The SAXS .dat files and derived models have been deposited in the Small Angle Scattering Biological Data Bank^66^. Accession numbers SASD2T2 (pH 5.6), SASD2U2 (pH 7.4 with calcium), SASD2V2 (pH 5.6 with calcium), SASD2W2 (pH 7.4 with calcium and Aleu-GFP), SASD2X2 (pH 5.6 and Aleu-GFP).

## Statistical analysis

Regression analysis were performed using Byos v5.10 (Protein Metrics). The dose response data representing fraction unmodified as a function of exposure time were fitted to a first-order exponential function, y = e−kt, to determine the hydroxyl radical reactivity rate constant, k (s−1). We filtered for high-quality data, retaining only cases in which the R2 of the exponential fit to the XFMS dose-response plot exceeded 0.9. Error estimates represent the standard error of the curve fit

## Acknowledgments

We would like to thank Ombretta Foresti for the initial cloning of VSR1 and help with the characterization of Aleu-GFP in plant cells. This work was supported by the Leverhulme Trust grant nr F/10 105/E. Further acknowledgements should be made to the Oxford protein production facility (OPPF) for the generation of the pOPIN-HBM-VSR1 vector and the initial purification of VSRs. We thank Louise Bird for her initial contribution while working at the OPPF, Maren Thompson for support and discussion. RO and JN were funded by the OPPF-UK grant MR/K018779/1. The authors wish to thank the staff of beamlines I04, I23, I24 and B21 and to the Membrane Protein Laboratory (MPL) for access to their facilities and for their expert support and assistance. The MPL is funded by Wellcome (223727/Z/21/Z). In particular, Nathan Cowieson and Nikul Katsuaki for discussion and support with collection of data on B21. Financial support was further provided by Instruct-ERIC (PID: 22309) and benefited from access to Diamond Light source Ltd, Harwell Science and Innovation Campus, Didcot, Oxfordshire, OX11 0DE, an Instruct-ERIC center. XFMS was supported in part by National Institute of Health grants P30 GM124169 and 1R01GM126218, and Laboratory Directed Research and Development funding. Work conducted at the Advanced Light Source and the Molecular Foundry was supported by the Office of Science of the U.S. DOE under contract DE-AC02-05CH1123.

RJB was supported by FAPESP (Process numbers 2016/24191-8 and 2017/13485-3) and CNPq (Process numbers 166593/2022-2) and MKE is supported by doctoral grant from Leeds Beckett University. The work was also supported by PID2021-128751NB-I00 (MICINN/AEI/FEDER/UE) to IU. RO and JN were funded by the OPPF-UK grant MR/K018779/1.

## Author Contributions

THW purified VSR1-Nt and Aleu-GFP, performed SEC-SAXS experiments including data analysis and modelling, conducted XFMS experiments and interpretation, revised manuscript. RJP purified VSR1, designed the experiments that led to the crystallisation of the cargo loaded form, collection of the crystallographic data and calcium phase information, jointly solved and analysed the 2.97Å cargo loaded form and solved the 1.87Å cargo loaded form, revised manuscript; RJB developed algorithm, jointly solved structure, performed modelling, analyzed structural results and revised manuscript; MKE cloned and purified Aleu-GFP in expression vector, optimised expression and purified VSR1-Nt for 2.97Å cargo-loaded form. NS performed crystallization trials for the crystal structure of the VSR1-Nt unloaded form. JA designed and performed initial cloning and expression assays. MP and PF performed the SEC-MALS experiments. LGK supported XFMS data acquisition and conducted the LC-MS/MS analysis and interpretation. SG led the XFMS data collection and contributed with interpretation. KEO supported data collection on I23 and data analysis of calcium sites; JB expressed and purified Aleu-GFP and VSR1-Nt (1.87Å cargo loaded form); AJLL performed modelling; MRMF designed modelling experiments and analyzed results; JD supported constructs and results analysis. RO supervised initial VSR cloning and expression assays; CYR supported XFMS data acquisition and analysis, AG analyzed results and revised manuscript, VP supervised research, designed experiments, analyzed results, revised manuscript; IU designed algorithm, analyzed results and revised manuscript, AQ supervised research and analysed data (cargo loaded),revised manuscript, CDML acquired funding, performed and coordinated experiments, supervised research, designed experiments, analyzed results, wrote and revised manuscript.

## SUPPLEMENTAL Material

**Supplemental Table 1.**
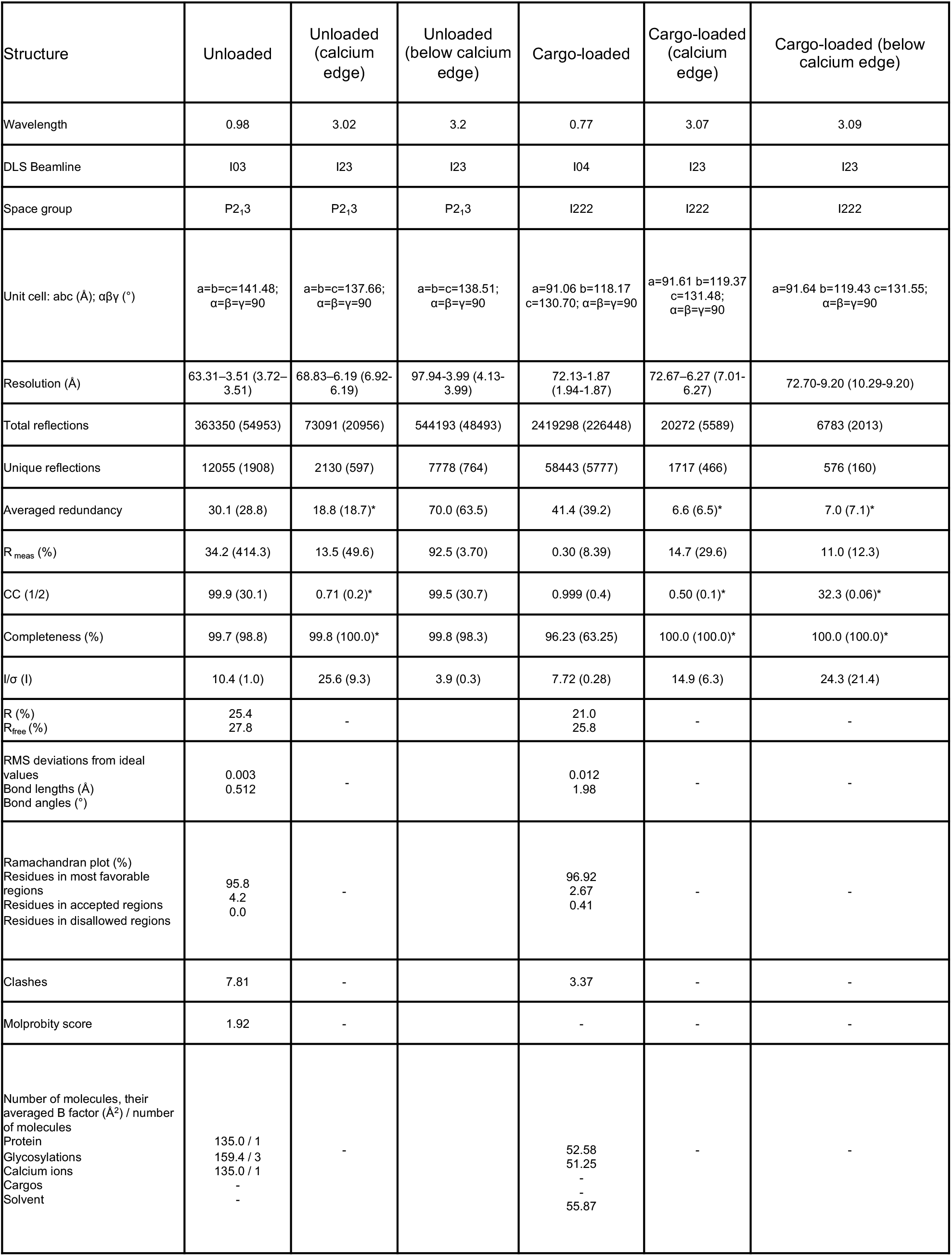
Diffraction data collection refinement statistics ^0^ RMS root-mean-square. * refers to anomalous data.

| Structure | Unloaded | Unloaded (calcium edge) | Unloaded (below calcium edge) | Cargo-loaded | Cargo-loaded (calcium edge) | Cargo-loaded (below calcium edge) |
|---|---|---|---|---|---|---|
| Wavelength | 0.98 | 3.02 | 3.2 | 0.77 | 3.07 | 3.09 |
| DLS Beamline | I03 | I23 | I23 | I04 | I23 | I23 |
| Space group | P2<sub>1</sub>3 | P2<sub>1</sub>3 | P2<sub>1</sub>3 | I222 | I222 | I222 |
| Unit cell: abc (Å); αβγ (°) | a=b=c=141.48; α=β=γ=90 | a=b=c=137.66; α=β=γ=90 | a=b=c=138.51; α=β=γ=90 | a=91.06 b=118.17 c=130.70; α=β=γ=90 | a=91.61 b=119.37 c=131.48; α=β=γ=90 | a=91.64 b=119.43 c=131.55;<br>α=β=γ=90 |
| Resolution (Å) | 63.31–3.51 (3.72–3.51) | 68.83–6.19 (6.92-6.19) | 97.94-3.99 (4.13-3.99) | 72.13-1.87 (1.94-1.87) | 72.67–6.27 (7.01-6.27) | 72.70-9.20 (10.29-9.20) |
| Total reflections | 363350 (54953) | 73091 (20956) | 544193 (48493) | 2419298 (226448) | 20272 (5589) | 6783 (2013) |
| Unique reflections | 12055 (1908) | 2130 (597) | 7778 (764) | 58443 (5777) | 1717 (466) | 576 (160) |
| Averaged redundancy | 30.1 (28.8) | 18.8 (18.7)* | 70.0 (63.5) | 41.4 (39.2) | 6.6 (6.5)* | 7.0 (7.1)* |
| R<sub>meas</sub> (%) | 34.2 (414.3) | 13.5 (49.6) | 92.5 (3.70) | 0.30 (8.39) | 14.7 (29.6) | 11.0 (12.3) |
| CC (1/2) | 99.9 (30.1) | 0.71 (0.2)* | 99.5 (30.7) | 0.999 (0.4) | 0.50 (0.1)* | 32.3 (0.06)* |
| Completeness (%) | 99.7 (98.8) | 99.8 (100.0)* | 99.8 (98.3) | 96.23 (63.25) | 100.0 (100.0)* | 100.0 (100.0)* |
| I/σ (I) | 10.4 (1.0) | 25.6 (9.3) | 3.9 (0.3) | 7.72 (0.28) | 14.9 (6.3) | 24.3 (21.4) |
| R (%)<br>R<sub>free</sub> (%) | 25.4<br>27.8 | - |  | 21.0<br>25.8 | - | - |
| RMS deviations from ideal values<br>Bond lengths (Å)<br>Bond angles (°) | 0.003<br>0.512 | - |  | 0.012<br>1.98 | - | - |
| Ramachandran plot (%)<br>Residues in most favorable regions<br>Residues in accepted regions<br>Residues in disallowed regions | 95.8<br>4.2<br>0.0 | - |  | 96.92<br>2.67<br>0.41 | - | - |
| Clashes | 7.81 | - |  | 3.37 | - | - |
| Molprobity score | 1.92 | - |  | - | - | - |
| Number of molecules, their averaged B factor (Å<sup>2</sup>) / number of molecules<br>Protein<br>Glycosylations<br>Calcium ions<br>Cargos<br>Solvent | 135.0 / 1<br>159.4 / 3<br>135.0 / 1<br>-<br>- | - |  | 52.58<br>51.25<br>-<br>-<br>55.87 | - | - |

**Supplemental Table 2.**
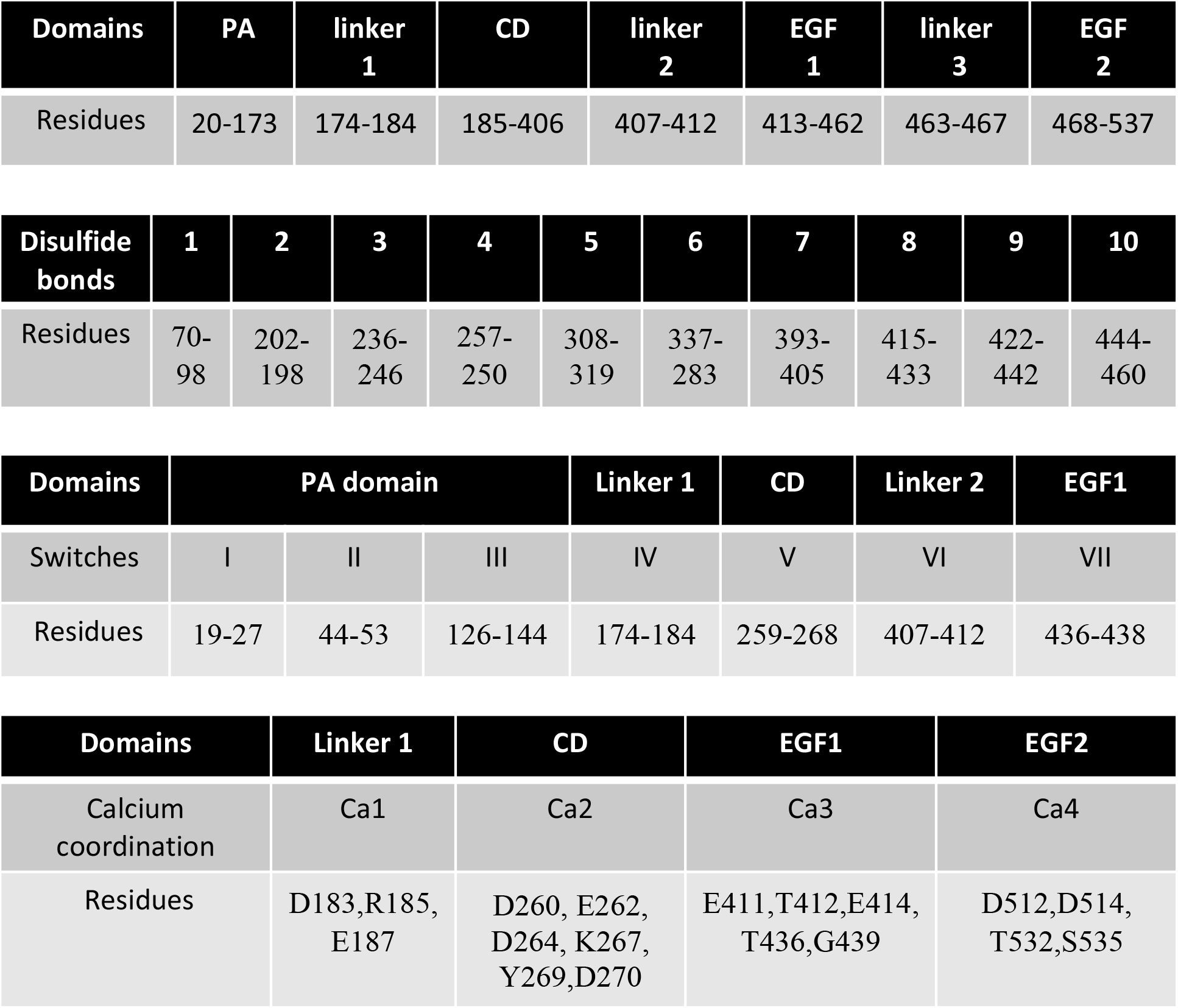
Structural properties of VSR1-N_t_ Describing domains, disulfide bonds, switches and calcium coordination.

**Supplemental Table 3.**
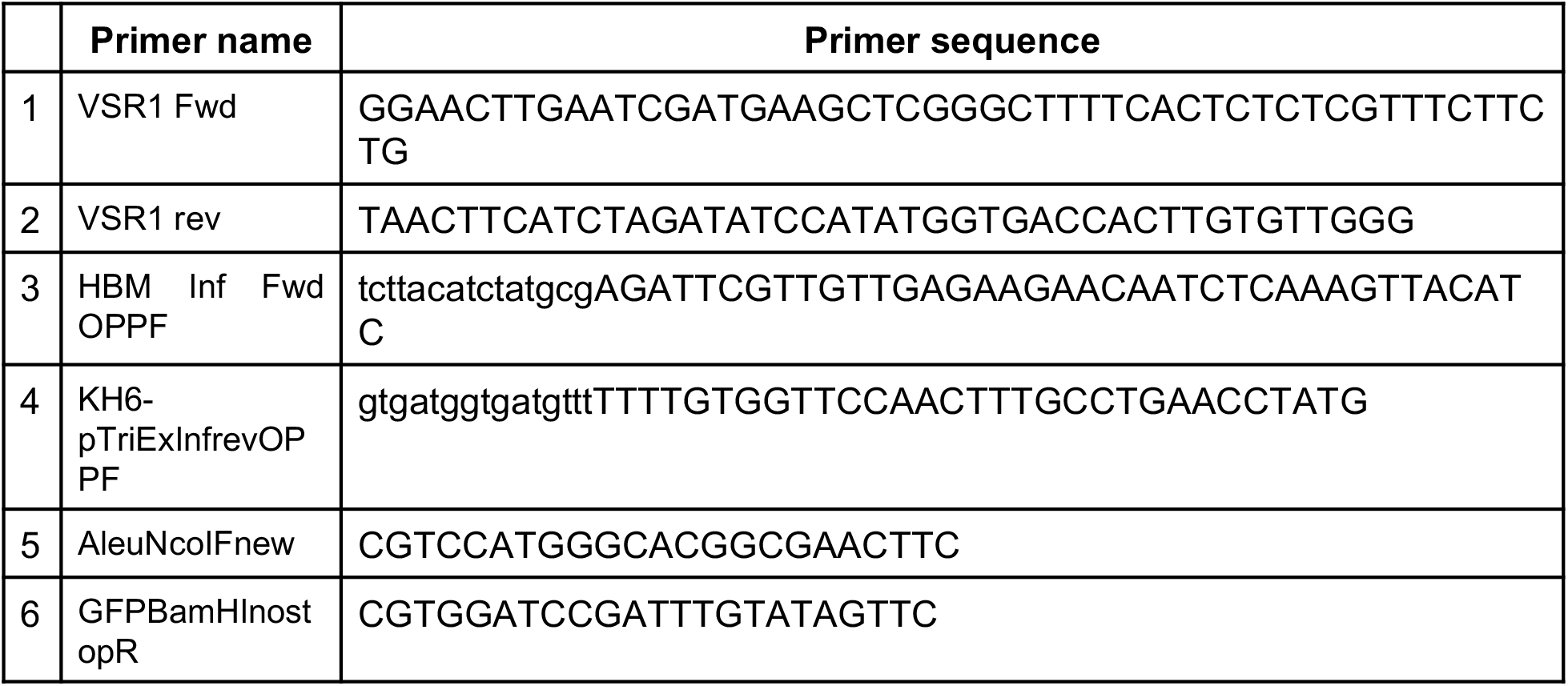
Primers of VSR1 and AleuGFP used in this study.

**Supplemental Table 4.** Anomalous peak heights (σ) detected by ANODE from anomalous difference Fourier maps. Only peaks above 4.5 σ threshold are shown.

| Sites | **3.07 Å** | **3.09 Å** |
|---|---|---|
| **Ca3** | 13.95 |  |
| **Ca2** | 13.40 |  |
| **Ca1** | 12.17 |  |
| **Ca4** | 6.51 |  |
| **C498/SG** | 6.30 | 6.41 |
| **Solvent** | 4.48 |  |
| **C250-C257/SGs** |  | 5.61 |
| **M310/SD** |  | 5.51 |
| **C422-C442/SGs** |  | 4.84 |
| **C42/SG** |  | 4.55 |

**Supplemental Table 5.**
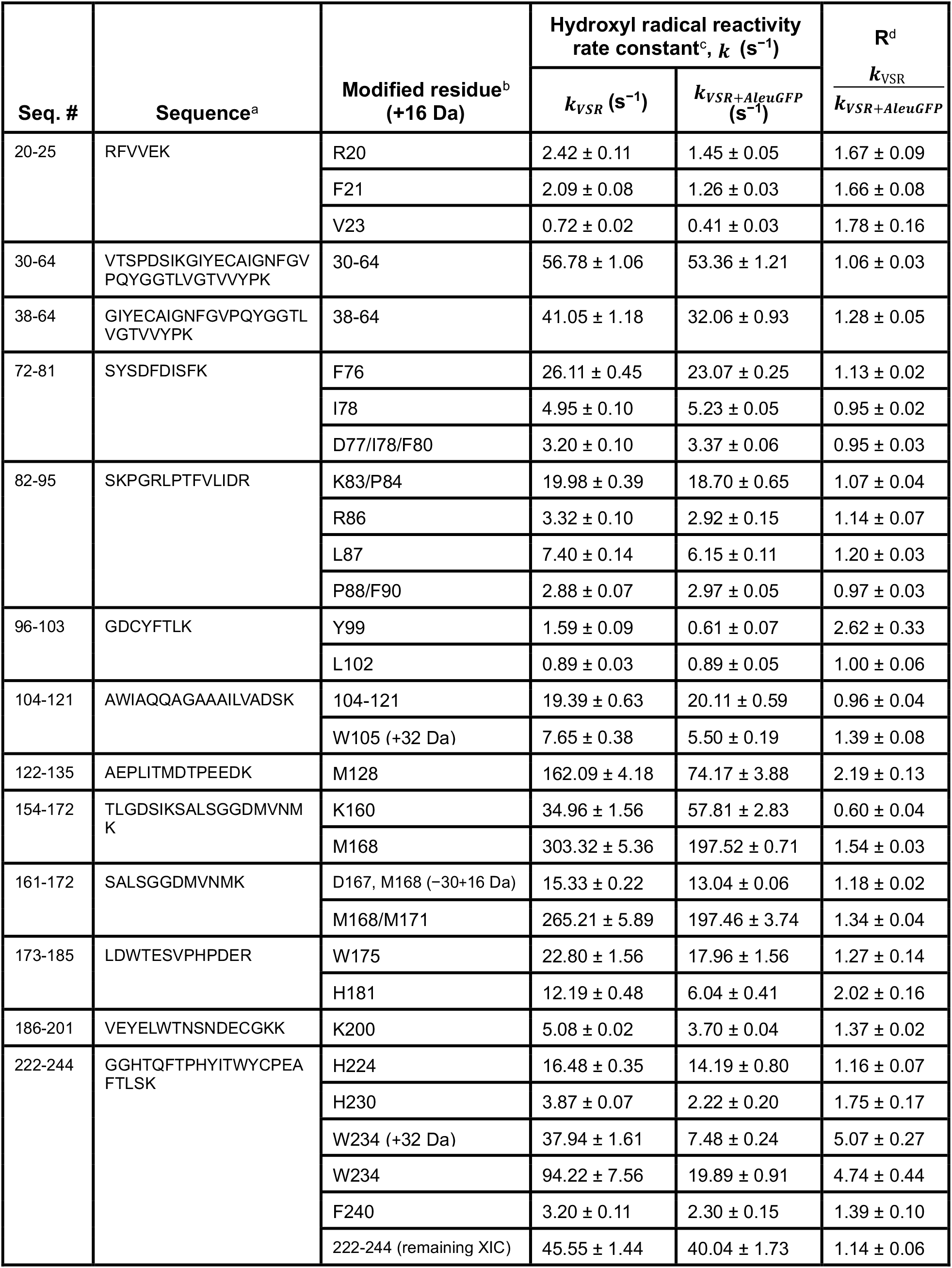
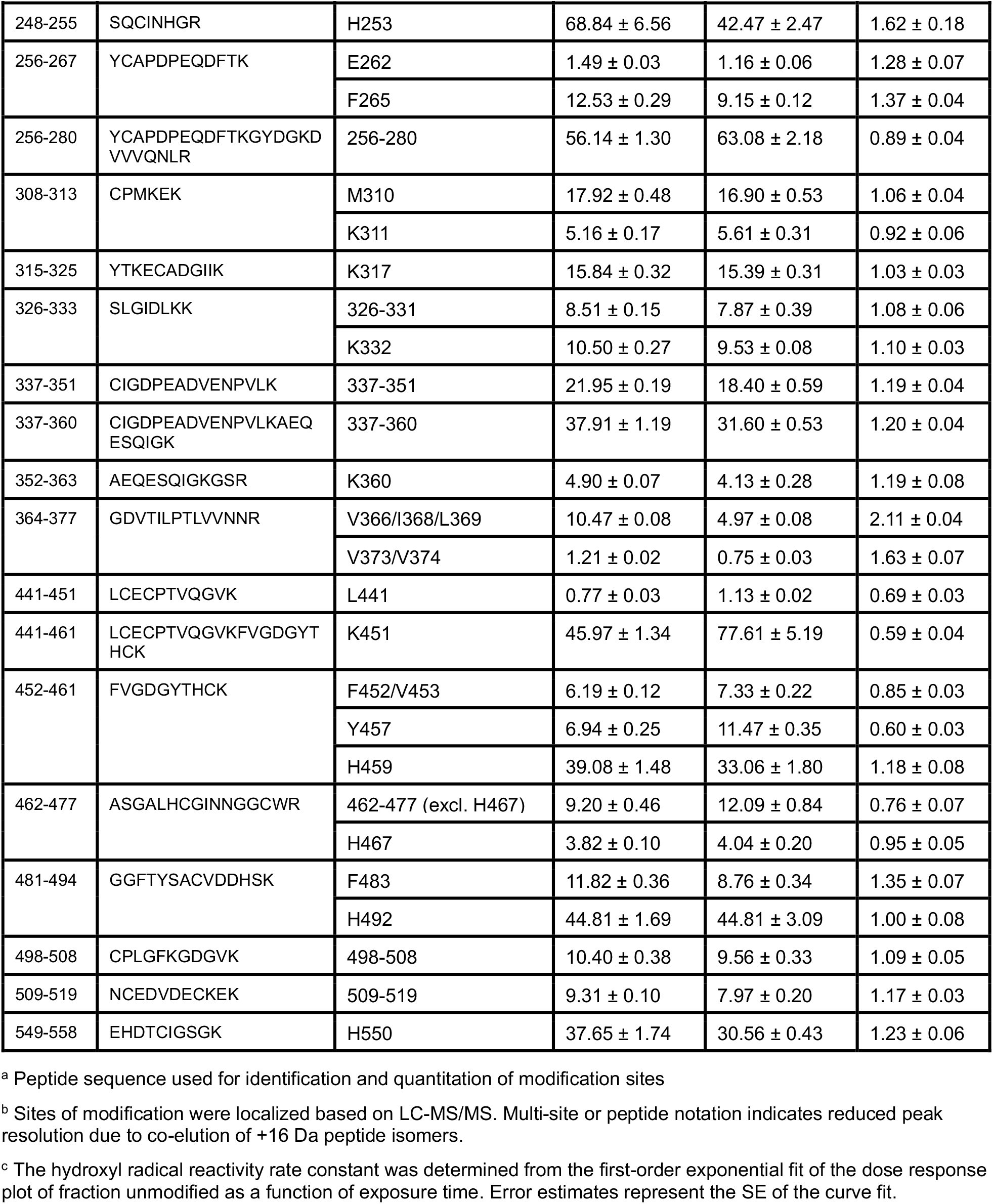
XFMS hydroxyl radical reactivity rate constants Supplementary Figure 1A.

| Seq. # | Sequence<sup>a</sup> | Modified residue<sup>b</sup> (+16 Da) | Hydroxyl radical reactivity rate constant<sup>c</sup>, *k* (s<sup>−1</sup>) |  | R<sup>d</sup> |
|---|---|---|---|---|---|
|  |  |  | *k<sub>VSR</sub>* (s<sup>−1</sup>) | *k<sub>VSR+AleuGFP</sub>* (s<sup>−1</sup>) | $\frac{k_{\mathrm{VSR}}}{k_{VSR+AleuGFP}}$ |
| 20-25 | RFVVEK | R20 | 2.42 ± 0.11 | 1.45 ± 0.05 | 1.67 ± 0.09 |
|  |  | F21 | 2.09 ± 0.08 | 1.26 ± 0.03 | 1.66 ± 0.08 |
|  |  | V23 | 0.72 ± 0.02 | 0.41 ± 0.03 | 1.78 ± 0.16 |
| 30-64 | VTSPDSIKGIYECAIGNFGVPQYGGTLVGTVVYPK | 30-64 | 56.78 ± 1.06 | 53.36 ± 1.21 | 1.06 ± 0.03 |
| 38-64 | GIYECAIGNFGVPQYGGTLVGTVVYPK | 38-64 | 41.05 ± 1.18 | 32.06 ± 0.93 | 1.28 ± 0.05 |
| 72-81 | SYSDFDISFK | F76 | 26.11 ± 0.45 | 23.07 ± 0.25 | 1.13 ± 0.02 |
|  |  | I78 | 4.95 ± 0.10 | 5.23 ± 0.05 | 0.95 ± 0.02 |
|  |  | D77/I78/F80 | 3.20 ± 0.10 | 3.37 ± 0.06 | 0.95 ± 0.03 |
| 82-95 | SKPGRLPTFVLIDR | K83/P84 | 19.98 ± 0.39 | 18.70 ± 0.65 | 1.07 ± 0.04 |
|  |  | R86 | 3.32 ± 0.10 | 2.92 ± 0.15 | 1.14 ± 0.07 |
|  |  | L87 | 7.40 ± 0.14 | 6.15 ± 0.11 | 1.20 ± 0.03 |
|  |  | P88/F90 | 2.88 ± 0.07 | 2.97 ± 0.05 | 0.97 ± 0.03 |
| 96-103 | GDCYFTLK | Y99 | 1.59 ± 0.09 | 0.61 ± 0.07 | 2.62 ± 0.33 |
|  |  | L102 | 0.89 ± 0.03 | 0.89 ± 0.05 | 1.00 ± 0.06 |
| 104-121 | AWIAQQAGAAAILVADSK | 104-121 | 19.39 ± 0.63 | 20.11 ± 0.59 | 0.96 ± 0.04 |
|  |  | W105 (+32 Da) | 7.65 ± 0.38 | 5.50 ± 0.19 | 1.39 ± 0.08 |
| 122-135 | AEPLITMDTPEEDK | M128 | 162.09 ± 4.18 | 74.17 ± 3.88 | 2.19 ± 0.13 |
| 154-172 | TLGDSIKSALSGGDMVNMK | K160 | 34.96 ± 1.56 | 57.81 ± 2.83 | 0.60 ± 0.04 |
|  |  | M168 | 303.32 ± 5.36 | 197.52 ± 0.71 | 1.54 ± 0.03 |
| 161-172 | SALSGGDMVNMK | D167, M168 (−30+16 Da) | 15.33 ± 0.22 | 13.04 ± 0.06 | 1.18 ± 0.02 |
|  |  | M168/M171 | 265.21 ± 5.89 | 197.46 ± 3.74 | 1.34 ± 0.04 |
| 173-185 | LDWTESVPHPDER | W175 | 22.80 ± 1.56 | 17.96 ± 1.56 | 1.27 ± 0.14 |
|  |  | H181 | 12.19 ± 0.48 | 6.04 ± 0.41 | 2.02 ± 0.16 |
| 186-201 | VEYELWTNSNDECGKK | K200 | 5.08 ± 0.02 | 3.70 ± 0.04 | 1.37 ± 0.02 |
| 222-244 | GGHTQFTPHYITWYCPEAFTLSK | H224 | 16.48 ± 0.35 | 14.19 ± 0.80 | 1.16 ± 0.07 |
|  |  | H230 | 3.87 ± 0.07 | 2.22 ± 0.20 | 1.75 ± 0.17 |
|  |  | W234 (+32 Da) | 37.94 ± 1.61 | 7.48 ± 0.24 | 5.07 ± 0.27 |
|  |  | W234 | 94.22 ± 7.56 | 19.89 ± 0.91 | 4.74 ± 0.44 |
|  |  | F240 | 3.20 ± 0.11 | 2.30 ± 0.15 | 1.39 ± 0.10 |
|  |  | 222-244 (remaining XIC) | 45.55 ± 1.44 | 40.04 ± 1.73 | 1.14 ± 0.06 |
| 248-255 | SQCINHGR | H253 | 68.84 ± 6.56 | 42.47 ± 2.47 | 1.62 ± 0.18 |
| 256-267 | YCAPDPEQDFTK | E262 | 1.49 ± 0.03 | 1.16 ± 0.06 | 1.28 ± 0.07 |
|  |  | F265 | 12.53 ± 0.29 | 9.15 ± 0.12 | 1.37 ± 0.04 |
| 256-280 | YCAPDPEQDFTKGYDGKDVVVQNLR | 256-280 | 56.14 ± 1.30 | 63.08 ± 2.18 | 0.89 ± 0.04 |
| 308-313 | CPMKEK | M310 | 17.92 ± 0.48 | 16.90 ± 0.53 | 1.06 ± 0.04 |
|  |  | K311 | 5.16 ± 0.17 | 5.61 ± 0.31 | 0.92 ± 0.06 |
| 315-325 | YTKECADGIIK | K317 | 15.84 ± 0.32 | 15.39 ± 0.31 | 1.03 ± 0.03 |
| 326-333 | SLGIDLKK | 326-331 | 8.51 ± 0.15 | 7.87 ± 0.39 | 1.08 ± 0.06 |
|  |  | K332 | 10.50 ± 0.27 | 9.53 ± 0.08 | 1.10 ± 0.03 |
| 337-351 | CIGDPEADVENPVLK | 337-351 | 21.95 ± 0.19 | 18.40 ± 0.59 | 1.19 ± 0.04 |
| 337-360 | CIGDPEADVENPVLKAEQESQIGK | 337-360 | 37.91 ± 1.19 | 31.60 ± 0.53 | 1.20 ± 0.04 |
| 352-363 | AEQESQIGKGSR | K360 | 4.90 ± 0.07 | 4.13 ± 0.28 | 1.19 ± 0.08 |
| 364-377 | GDVTILPTLVVNNR | V366/I368/L369 | 10.47 ± 0.08 | 4.97 ± 0.08 | 2.11 ± 0.04 |
|  |  | V373/V374 | 1.21 ± 0.02 | 0.75 ± 0.03 | 1.63 ± 0.07 |
| 441-451 | LCECPTVQGVK | L441 | 0.77 ± 0.03 | 1.13 ± 0.02 | 0.69 ± 0.03 |
| 441-461 | LCECPTVQGVKFVGDGYTHCK | K451 | 45.97 ± 1.34 | 77.61 ± 5.19 | 0.59 ± 0.04 |
| 452-461 | FVGDGYTHCK | F452/V453 | 6.19 ± 0.12 | 7.33 ± 0.22 | 0.85 ± 0.03 |
|  |  | Y457 | 6.94 ± 0.25 | 11.47 ± 0.35 | 0.60 ± 0.03 |
|  |  | H459 | 39.08 ± 1.48 | 33.06 ± 1.80 | 1.18 ± 0.08 |
| 462-477 | ASGALHCGINNGGCWR | 462-477 (excl. H467) | 9.20 ± 0.46 | 12.09 ± 0.84 | 0.76 ± 0.07 |
|  |  | H467 | 3.82 ± 0.10 | 4.04 ± 0.20 | 0.95 ± 0.05 |
| 481-494 | GGFTYSACVDDHSK | F483 | 11.82 ± 0.36 | 8.76 ± 0.34 | 1.35 ± 0.07 |
|  |  | H492 | 44.81 ± 1.69 | 44.81 ± 3.09 | 1.00 ± 0.08 |
| 498-508 | CPLGFKGDGVK | 498-508 | 10.40 ± 0.38 | 9.56 ± 0.33 | 1.09 ± 0.05 |
| 509-519 | NCEDVDECKEK | 509-519 | 9.31 ± 0.10 | 7.97 ± 0.20 | 1.17 ± 0.03 |
| 549-558 | EHDTCIGSGK | H550 | 37.65 ± 1.74 | 30.56 ± 0.43 | 1.23 ± 0.06 |

<sup>a</sup> Peptide sequence used for identification and quantitation of modification sites
<sup>b</sup> Sites of modification were localized based on LC-MS/MS. Multi-site or peptide notation indicates reduced peak resolution due to co-elution of +16 Da peptide isomers.
<sup>c</sup> The hydroxyl radical reactivity rate constant was determined from the first-order exponential fit of the dose response plot of fraction unmodified as a function of exposure time. Error estimates represent the SE of the curve fit.

---

**Supplemental Figure 1.**
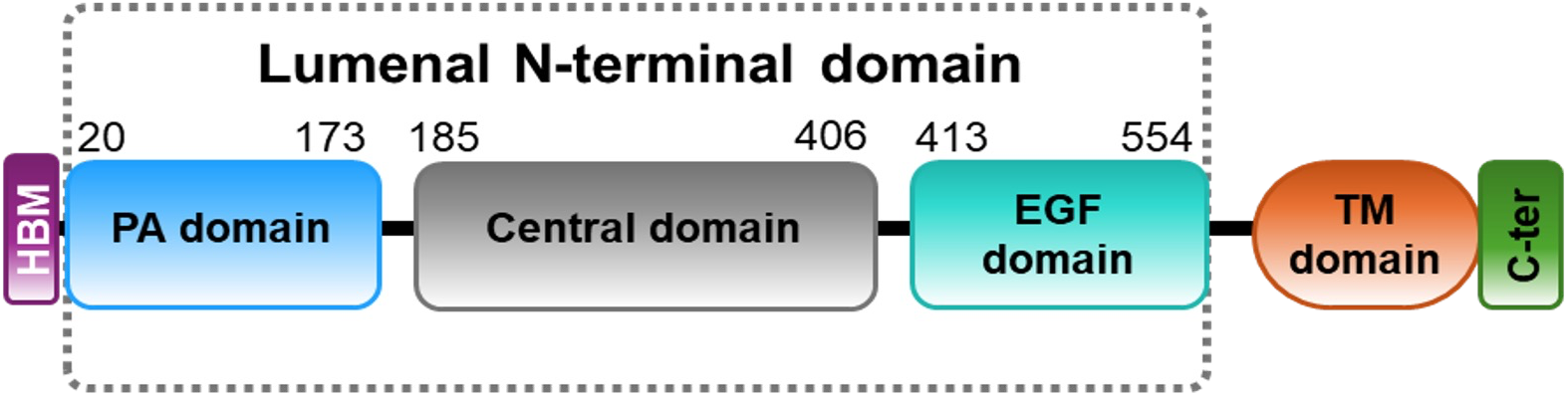
Purified N-terminal VSR1 lumenal domain. **A.** Schematic of VSR1 protein with the different domains. The lumenal N-terminal domain used in this study comprised the three N-terminal domains (PA, Central and EGF repeats) fused to Honey Bee Melittin secretion signal (HBM) and Histidine tag.

**Supplemental Figure 2.**
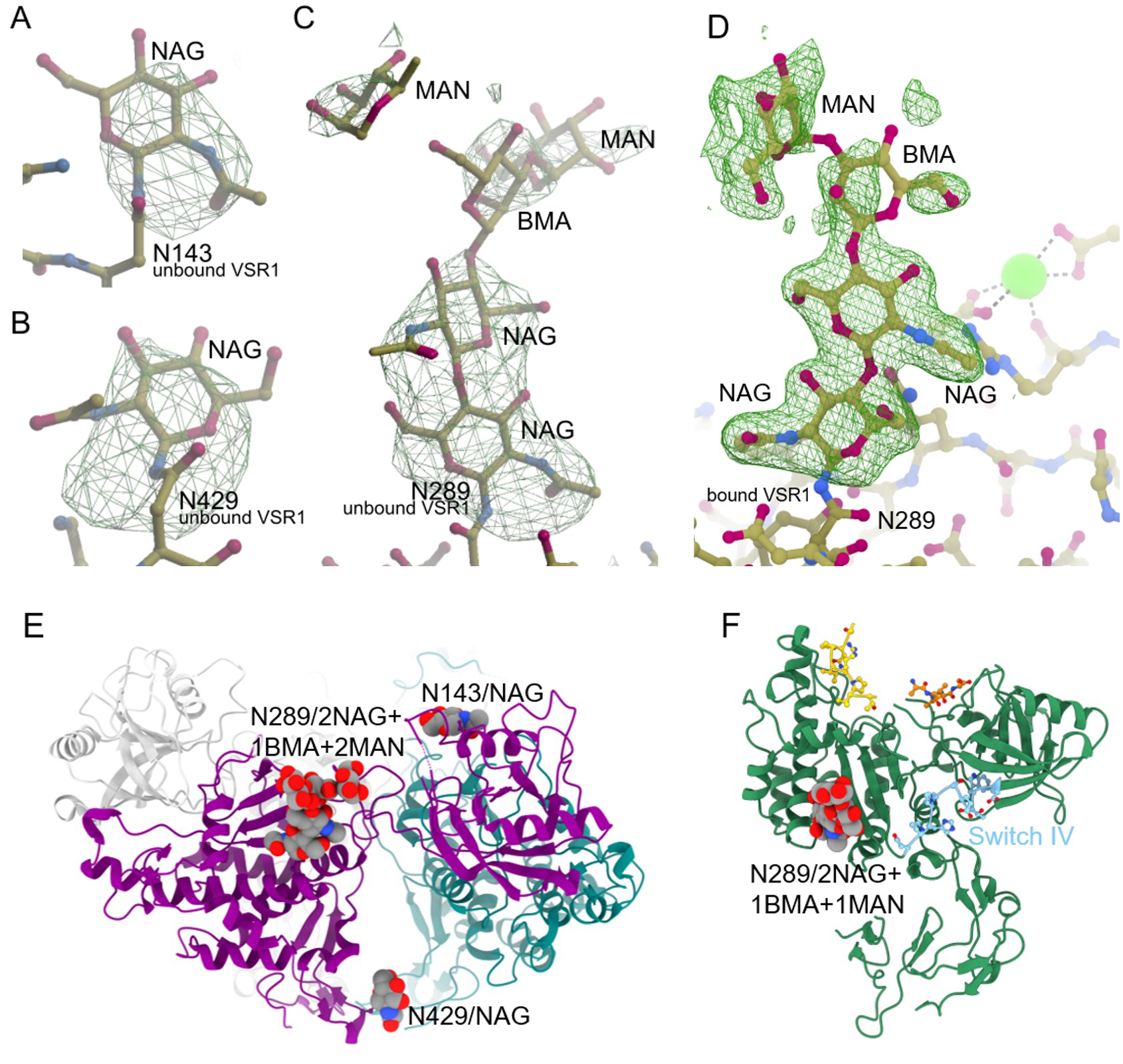
Glycosylations with their omit electron density (3 σ) and their localisation in both unloaded and cargo-loaded VSR1-N_t._ A. The unbound VSR1-N_t_ structure includes a 2-acetamido-2-deoxy-beta-D-glucopyranose (NAG) bound to asparagine 143 (N143). B. Binding to N429/NAG C. N289/2NAG bound to β-D-mannopyranose (BMA) bound to an α-D-mannopyranose (MAN). D. The cargo-loaded VSR1-N_t_ has N289/NAG/NAG/R185 in a site close to coordinated calcium ion. E. The three glycosylations locations within the trimer structure. F. The N289/2NAG in the cargo-loaded monomer showing its location close to switch IV.

**Supplemental Figure 3.**
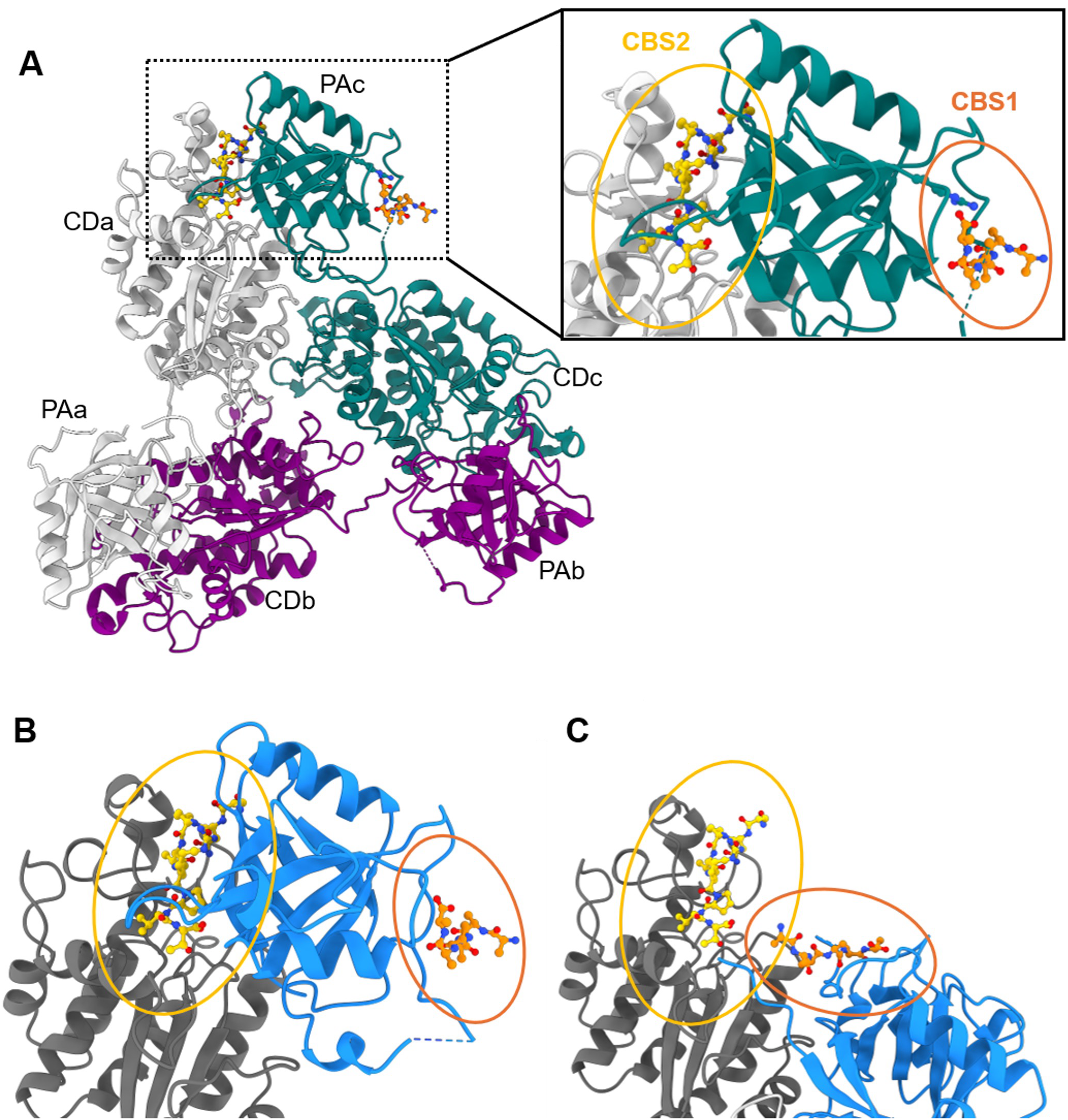
The trimer occludes the exposition of cargo binding site 2 and reorganised. A. The location of the two peptides is artificially shown within the trimer to demonstrate that cargo binding sites 2 is not accessible but that cargo bind site 1 is. B. Zoom into the trimer (left, with blue PA domain of unloaded structure a and grey Central domain of unloaded structure) C. Right, for comparison with the cargo-loaded structure.

**Supplemental Figure 4.**
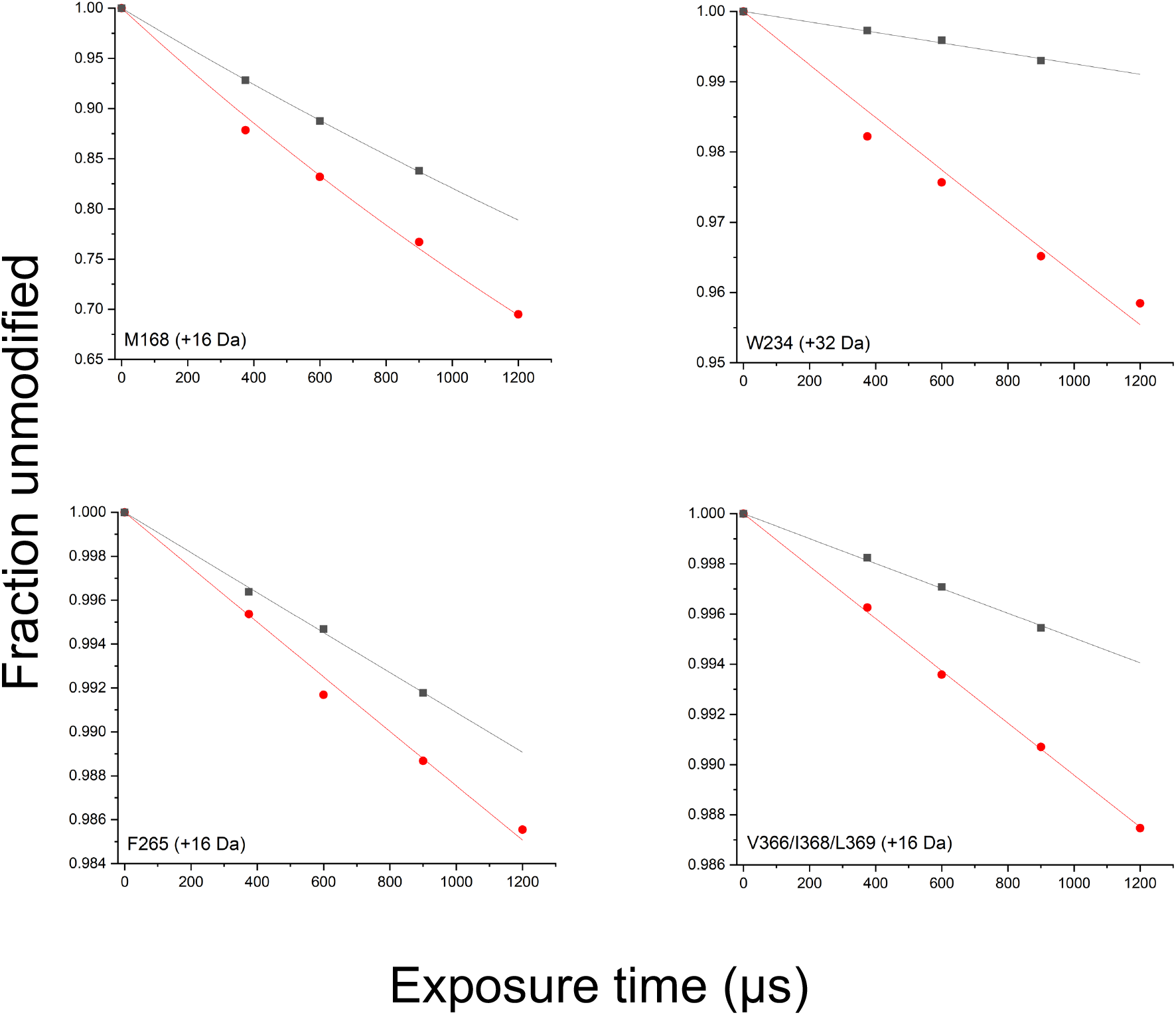
Representative XFMS dose response plots. Fraction unmodified as a function of exposure time for VSR1-N_t_ (●) and VSR1-N_t_+Aleu-GFP (▪).The solid line represents a first-order exponential fit to determine the hydroxyl radical reactivity rate constant, *k* (s^−1^).

**Supplemental Figure 5.**
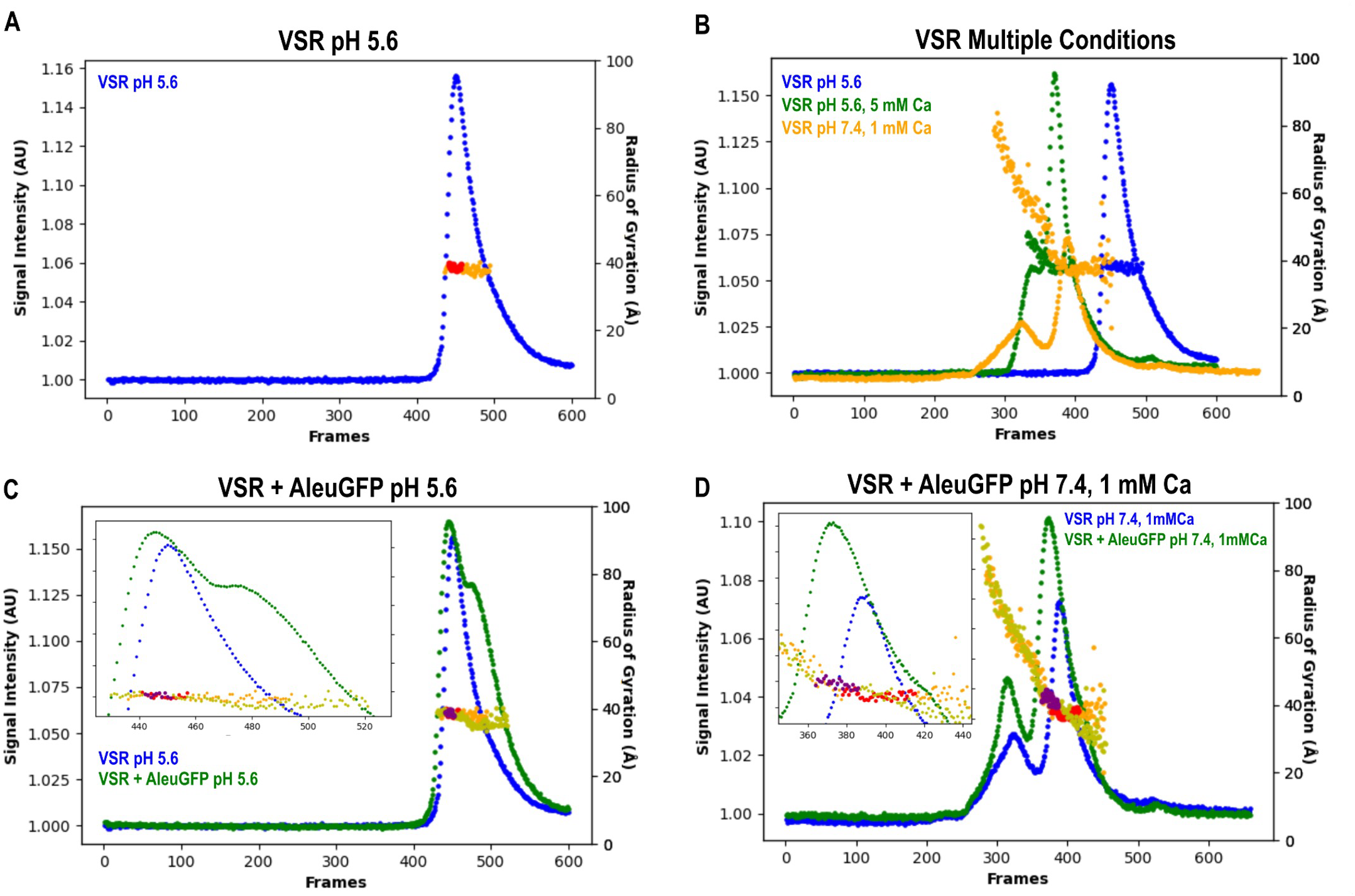
SEC-SAXS Elution profiles of VSR1-Nt. **A)** SEC-SAXS profile (blue) at pH 5.6, with radii of gyration (orange). Frames merged for SAXS analysis are in red. **B)** SEC-SAXS profiles and radii of gyration at pH 5.6, pH 5.6 5 mM Ca and pH 7.4 1 mM Ca. **C)** SEC-SAXS profile at pH 5.6, coloured as in **(A),** and with 1:1 Aleu-GFP (green) with radii of gyration (olive yellow). Frames merged for SAXS analysis are in red (alone) and purple (+ Aleu-GFP). Inset: Zoom in over main peak. **D)** SEC-SAXS profile at pH 7.4 1 mM Ca, coloured as in **(A),** and with 1:1 Aleu-GFP (green) with radii of gyration (olive yellow). Frames merged for SAXS analysis are in red (alone) and purple (+ Aleu-GFP). Inset: Zoom in over main peak.

## Supplemental detailed Materials and Methods

### Constructs

The Vacuolar sorting receptor VSR1 (AT3G52850) was amplified from *Arabidopsis thaliana* genomic DNA using the primers 1 and 2 and cloned in a pUC vector to form pOF128. The lumenal region of the VSR1 gene (VSR1-N) was further amplified with primers 3 and 4 to insert into the pOPIN-HBM vector (Oxford Protein Production Facility, OPPF). The gene is flanked by the signal sequence honeybee melittin leader (HBM) sequence to direct the secretion of the VSR1 in the cell culture of Expi293 cells, and a C-terminal octahistidine tag followed by a T7 terminator.

Petunia Aleurain (Aleu) construct was obtained from the Roger laboratory ^67^. Aleu was amplified and cloned as a GFP fusion protein in a pUC vector to generate the pJA1 vector. Using primers 5 and 6, the Aleu-GFP cassette was then subcloned into the pCRBlunt to isolate the Aleu peptide [TANFADE**NPIR**QVVSDSFHELESAS] in frame with the GFP. The cassette was then transferred into the pET28b (Novagen) with a partial NcoI/complete BamHI digest, resulting in the pME4, where Aleu-GFP is in frame with a C-terminal hexahistidine tag. All constructs were sequenced at each stage of cloning to confirm identity. A list of primers is presented in **SI Appendix, Table 3**.

### Expression and purification of VSR and AleuGFP

The lumenal N-terminal region of VSR1 was expressed in Expi293 cells in the presence or absence of kifunensin 1 mg/ml following the published protocol ^42^. Briefly, the lumenal region fused to an HBM motif was expressed for 4 days in Expi293 cells (Thermofisher). The supernatant (containing the secreted HBM-VSR1) was collected by centrifugation then VSR1 purified using affinity chromatography followed by size exclusion chromatography. Fractions were pooled and VSR1 was resuspended in 20 mM Tris pH 7.4, 200 mM NaCl. Thirty ml of Expi cell culture resulted in a yield of 4.25 mg of VSR1. This was further concentrated to 15 mg/ml and used for further crystallization studies.

Aleu-GFP bacterial expression was performed as follows: AleuGFP was expressed in *Escherichia coli* BL21 Star (DE3) (ThermoFisher) by autoinduction at 37° C for 16 hours ^43^. Cells were collected by centrifugation at 4,000 g for 15 minutes. The obtained pellet was resuspended in equilibration buffer 1 (50 mM Tris-HCl, 200 mM NaCl, 5 mM imidazole pH 8.0). A final concentration of 0.5 mg/ml of lysozyme was added and incubated for 45 min at 4° C. The tube containing the solution was placed on ice and sonicated with a probe sonicator (Soniprep 150, MES) for 20 seconds at 10-second intervals until the turbidity was clear. The solution was then centrifuged for 10 min at 14,000 g (4° C) to remove unbroken cells and cell debris. The supernatant was collected and then incubated with Ni beads (prewashed in buffer 1) for 1 h at RT. The flowthrough was removed and the beads were then washed with 2 volumes of buffer 1 and then 2 volumes of buffer 2 (50 mM Tris, 200 mM NaCl, 15 mM imidazole pH 8.0). Finally, Aleu-GFP was eluted in 1 volume of elution buffer (50 mM Tris, 200 mM NaCl, 250 mM imidazole pH 8.0). The protein purification was confirmed on an SDS PAGE stained with Coomassie and by Western blot using 1:5000 anti-His-tag horseradish peroxidase-conjugated antibody (MAB050H, R&D Systems).

The Aleu-GFP was further purified with size exclusion chromatography via ÄKTA pure protein purification system (Cytiva) using Superdex 200 Increase 10/300 GL column (Cytiva) using 1.5 CV of 20 mM Tris pH 8.0, 200 mM NaCl. Fractions containing the fluorescent protein were pooled and the protein was concentrated with Vivaspin 20 MWCO 10,000 (Cytiva) to the desired concentration.

### Crystallographic determination of unbound VSR1 structure

Due to a lack of electron density, residues 129–135 were omitted in the crystallographic model of the unbound VSR1. Positive peaks in the difference map were found close to the side chains of Asn143, Asn289 and Asn429 in agreement with reported glycosylation sites ^20^. Omit electron density map supported a single 2-acetamido-2-deoxy-β-D-glucopyranose (NAG) for Asn143 and Asn429 glycosylation, while for Asn289 it was possible to include 2NAG-1BMA(β-D-mannopyranose)-2MAN(α-D-mannopyranose) (**SI Fig. 2**). Due to the limited data of this low-resolution crystal structure, parameters refined in phenix.refine ^54^ were reduced by using grouped Bfactors refinement. Due to weak electron density, BMA and MANs had their occupancy reduced to 0.75 and side chains of residues 20, 21, 22, 24, 25, 86, 93, 95, 99, 102, 121, 125, 126, 128, 139, 141, 153, 155, 163, 168, 176, 177, 181, 189, 197, 198, 207, 234, 238, 242, 250, 257, 263, 264, 265, 267, 270, 283, 308, 311, 317, 318, 325, 332, 337, 342, 355, 360, 380, 384, 390, 393, 405, 415, 422, 433, 434, 438, 442, 444, 455, 457, 460 had Bfactors increased to a factor of 1.5 of main chain Bfactor. Probably due to radiation damage, negative peaks appeared in sulfurs of cysteines 70, 98, 202, 236, 246 and 319, therefore we included its side chain reduced form with a half occupancy. The final model had R/Rfreeof 25.4/27.8%, 95.8% of Ramachandran favored residues and no outliers, 1.9% poor rotamers and no Cβ outliers, a clashscore of 7.8 and a Molprobity score of 1.92, which is in the 100^th^ best percentile of all structures elucidated from datasets of similar resolution (3.25–9.93 Å) ^68^. The quaternary assembly of VSR1 was predicted using PISA software ^69^ and interaction between monomers was calculated with PLIP ^70^. Figures were generated using Coot ^71^ Chimera X 1.11.1^72^, and PyMOL (The PyMOL Molecular Graphics System, Version 1.8.4.0, Schrödinger, LLC., http://www.pymol.org/).

### Crystallographic determination of cargo-bound VSR structure

Due to lack of electron density, residues 133-137 were omitted in the crystallographic model of bound VSR1. Positive peaks in the difference map were found close to the side chain of Asn289 in agreement with reported glycosylation sites^20^. Electron density corresponding to peptide binding was observed in CBS1; however, this density was left unmodelled. Although a polyalanine trace could be built and the C-terminal carboxyl group located, the density was incompatible with the proline-containing ssVSD sequence, resulting in insufficient confidence in peptide placement and register assignment. By comparison, the density in CBS2 enabled modelling of 8 of the 10 ssVSD peptide residues. The final model had R/Rfreeof 21.0/25.8 %, 96.9 % of Ramachandran favored residues and 0.4 % outliers, 1.2 % poor rotamers and a clashscore of 3.37. Figures were generated using Chimera X 1.11.1^72^.

### XFMS Peptide Digestion, LC-MS/MS and Data Analysis

Samples were irradiated at the Advanced Light Source (ALS) beamline 3.3.1 (Lawrence Berkeley National Laboratory), which delivers a 3–12 keV broadband X-ray beam from a bending-magnet source with a high-flux, focused beam^64^. Using an X-ray slit, the beam was configured to produce a full width at half maximum of 640 μm along the vertical axis and 250 μm along the horizontal axis. Sample exposure was carried out by loading the sample into a 1 mL gas-tight Luer-lock Hamilton glass syringe, which was positioned in a syringe pump that flowed the sample through a 200 μm ID polymicro-coated fused silica capillary (Molex®) at various speeds along the vertical axis of the beam. An aliquot of each free VSR1-Nt sample was used for automated Alexa 488 fluorescence dose-response analysis to assess hydroxyl radical dose variability between samples^73^. X-ray exposure times were 375, 600, 900, and 1200 µs. The leftover unexposed sample in the syringe was used for the 0 μs control. Irradiated samples were collected in LoBind® Eppendorf tubes containing L-methionine amide (10 mM final concentration) to quench any secondary reactions. All samples were stored at −80°C until further processing by bottom-up LC-MS/MS.

A 60 µL aliquot per exposure was reduced to ∼10 µL in a vacuum centrifuge. Samples were denatured and reduced at 37°C for 30 min using 6 M guanidine hydrochloride (GuHCl), 7.5 mM dithiothreitol (DTT); followed by dilution to 0.5 M GuHCl, 1 mM DTT using 50 mM ammonium bicarbonate. Trypsin/Lys-C protease mix (Promega) was added at a 1:20 protease to protein ratio (w/w) and samples were incubated at 37°C for 12 h. Post-digestion sample reduction and acidification were performed using 1 mM TCEP and 0.5% formic acid. Peptides were desalted on C18 spin columns (Pierce, Thermo Fisher Scientific), dried in a vacuum centrifuge, and reconstituted in 25 µL 0.1% formic acid in water.

Peptides were analyzed using a Vanquish Flex UHPLC system coupled to an Orbitrap Exploris 480 mass spectrometer (Thermo Scientific). Separation was performed on a Thermo Scientific Hypersil GOLD Peptide C18 column (2.1 x 100 mm, 1.9 µm particle size, maintained at 55°C) at a flow rate of 0.400 mL/min with a mobile phase consisting of solvent A (0.1% formic acid in water) and solvent B (99.9% acetonitrile, 0.1% formic acid). The injection volume was 20 µL and peptides were eluted with the following gradient: initial condition was 2% solvent B which was increased to 10% over 1.5 min, then increased to 35% over 15 min followed by a ramp-up to 80% over 0.5 min and held at 80% solvent B for 1.5 min at a flow rate of 0.600 mL/min (flow was diverted to waste at 18 min). The column underwent a sawtooth wash before being equilibrated to 2% solvent B. The mass spectrometer was operated in positive mode with full scan parameters set to an Orbitrap resolution of 60,000; mass range 300–1200 m/z; RF lens at 50%; AGC target at 3.0e6; maximum injection time 60 ms. The top 10 most intense ions per MS scan were selected for HCD fragmentation with an intensity threshold of 5.0e3 and dynamic exclusion set at 2 s. Data-dependent MS2 scan parameters included an isolation window of 2 m/z; 30% normalized collision energy; Orbitrap resolution set to 15,000; AGC target at 1.0e5; maximum injection time 50 ms.

Peptide identification, localization of modification sites, quantitation, and regression analysis were performed using Byos v5.10 (Protein Metrics). The precursor mass tolerance was 10 ppm and the fragment mass tolerance was 20 ppm. Digestion parameters reflected the expected trypsin/Lys-C cleavage sites, allowing up to 2 missed cleavages. Variable oxidative modifications were specified for 18 amino acid side chains with mass shift search parameters of +14, +16, +32, and +48 Da. Identified tryptic peptides covered 100% of the VSR1-Nt protein sequence. Byos-assigned, site-specific modifications were validated manually using MS2 spectra. A series of hydroxyl radical doses, obtained by varying the sample exposure time to the X-ray beam, produced a range of oxidative modifications. Based on extracted ion chromatograms, the software automatically calculated fraction unmodified for each peptide as the ratio of the unmodified peptide peak area to the sum of the peak areas of the unmodified and modified peptides. The dose response data representing fraction unmodified as a function of exposure time were fitted to a first-order exponential function, y = e−kt, to determine the hydroxyl radical reactivity rate constant, k (s−1). We filtered for high-quality data, retaining only cases in which the R^2^ of the exponential fit to the XFMS dose-response plot exceeded 0.9. Several residues, such as Cys, Thr, Ser, Gln, Asn, and Gly, typically do not yield good dose-response plots due to complex secondary reactions with hydroxyl radicals or low intrinsic reactivity toward radicals. In comparison, the best XFMS data often derive from aromatic and hydrophobic residues.

